# Distinct molecular and functional properties of human induced-proprioceptor and low-threshold mechanoreceptor neurons

**DOI:** 10.1101/2024.05.12.593730

**Authors:** Amy J. Hulme, Rocio K. Finol-Urdaneta, Jeffrey R. McArthur, Nicholas R. Marzano, Simon Maksour, Amarinder Thind, Yang Guo, Dominic Kaul, Marnie Maddock, Oliver Friedrich, Boris Martinac, David J. Adams, Mirella Dottori

**Author notes:** Correspondence, Building 32, Northfields Avenue, University of Wollongong, Wollongong, 2522 NSW, Australia.

## Abstract

Sensing mechanical stimuli is crucial for the function of internal and external tissues, such as the skin and muscles. Much of our understanding of mechanosensory physiology relies on rodent studies, which may not directly translate to humans. To address the knowledge gap in human mechanosensation, we developed distinct populations of human mechanosensory neuronal subtypes from human pluripotent stem cells (hPSC). By inducing co-expression of NGN2/RUNX3 or NGN2/SHOX2 in hPSC-derived migrating neural crest cells we directed their specification to proprioceptor and low-threshold mechanoreceptor neuronal subtypes, respectively. The induced neurons exhibited transcriptional profiles consistent with mechanosensory neurons and displayed functional responses to mechanical stimuli, such as stretch and submicrometer probe indentation to the soma. Notably, each subtype displayed unique mechanical thresholds and desensitization properties akin to proprioceptors and low-threshold mechanoreceptors and both induced neuronal subtypes fired action potentials in response to minute mechanical stimuli, predominantly relying on PIEZO2 for mechanosensory function. Collectively, this study provides a foundational model for exploring human neuronal mechanosensory biology.

## Introduction

Mechanosensation, the mechanism by which mechanical stimuli is detected and converted into biochemical signals, is essential for everyday functions such as sitting, walking, and holding objects, as well as, detecting internal organ sensations, e.g., in the GI tract and bladder. Mechanosensory neurons, whose cell bodies reside within the dorsal root ganglia (DRG), are specialised neurons that detect mechanical stimuli and are broadly divided into two major classes: (i) proprioceptors (PNs) that detect mechanical signals such as joint position and muscle tension, and (ii) low threshold mechanoreceptors (LTMRs) which detect mechanical cues such as touch, hair deflection and vibration. Classification of mechanosensory neuron subtypes is multifactorial and includes their innervation of target tissues, transcriptional profiles of membrane receptors, ion channels, and transcription factors, and their distinct functional abilities to respond to and discriminate between varying types of mechanical stimuli and intensities ^1–4^. Most of our understanding of mechanosensory neuronal functions comes from rodent DRG neurons due to limited access to human primary DRG tissue. Yet, recent advances in single-cell technologies have begun to identify species-specific sub-classes of LTMRs and PNs ^5–7^. Critically, an increasing number of studies show inherent species differences between human and rodent DRGs, which questions the translation of rodent-based findings to humans ^8–11^.

Advances in human pluripotent stem cell (hPSC) biology have been a cornerstone for bridging this gap of knowledge in human neuroscience as differentiation methodologies are becoming increasingly optimised for generating specific neuronal populations. To this end, genetic engineering is growing as a preferable approach for consistently generating homogenous cell types ^12,13^. The basis of this methodology is to intrinsically regulate the expression of transcription factors that drive cell lineage specification. This strategy has been successfully utilised for generating mechanosensory neurons from hPSC, however, the transcription factors used include NEUROGENIN 2 (NGN2), NEUROGENIN 1 (NGN1) and/or BRN3A, all of which are expressed across multiple DRG sensory subtypes during development ^14–19^. Indeed, findings from our laboratory and others, demonstrate that NGN2-induced DRG sensory neurons generate a heterogeneous population of DRG subtypes ^17–19^, which confound comprehensive molecular and functional analyses to define specific subtypes in humans. We, therefore, propose that combining NGN2 expression with other transcription factors known to drive the specification of mechanosensory subclasses *in vivo* may be a strategy to generate similar mechanosensory populations from hPSC-derived progenitors. Based on developmental studies in rodent and chick ^20–25^, RUNX3 and SHOX2 were chosen as candidate transcription factors, to be co-expressed with NGN2 in hPSC-derived progenitors, as inducers of PNs and LTMRs differentiation, respectively.

In this study, we mimicked sensory neurogenesis via a multistep differentiation protocol, to generate neural crest cells, followed by the timely induced expression of NGN2 combined with either RUNX3 or SHOX2, to generate induced-proprioceptors (iPNs) and induced-LTMRs (iLTMRs), respectively. The iPNs and iLTMRs displayed expression and functional characteristics akin to native proprioceptors and LTMRs, respectively, and responded to different modes of mechanical stimulation. iPNs and iLTMRs exhibited exquisitely sensitive responses to mechanical stimulation and had distinct differences in the mechanical thresholds and desensitization properties reflective of distinct sensory specializations. Importantly, iPNs and iLTMRs fired action potentials in response to mechanical stimulation. Overall, our findings describe a unique platform for understanding the fundamental mechanisms that govern distinct responses of human mechanosensory neuronal subtypes to mechanical stimuli that give rise to the perception of touch and tension.

## Results

### Expression of NGN2 with RUNX3 or SHOX2 in hPSC-derived neural crest cells induces molecular profiles consistent with PN and LTMR neurons, respectively

We previously described a robust protocol for generating DRG-induced sensory neurons from hPSC via induced expression of NGN2 in neural crest progenitors ^18^. This method generates a mixed population of functional DRG sensory neurons that display functional expression of ion channels akin to DRG sensory neurons. Using this approach, hPSCs were initially differentiated into caudal neural progenitors, followed by neural crest cells, and further enriched for migrating neural crest cells (Figure 1A) ^18,26^. To guide the differentiation of these migrating neural crest cells towards populations of mechanosensory neuron subtypes, we transiently induced either *NGN2* and *RUNX3* or *NGN2* and *SHOX2* expression, via lentiviral vectors (Figure S1), respectively, and the cultures were then further matured to neurons (Figure 1A). Both resulting induced sensory neuronal cultures exhibited molecular profiles consistent with mechanosensory neurons (Figure 1). Bulk RNA sequencing unveiled elevated expression of transcripts associated with neurons such as *MAP2*, *TUBB3* (ß-III-TUBULIN), *RBFOX3* (NeuN), and sensory neuron-specific markers like *PRPH* (PERIPHERIN), *NEFH* (Neurofilament heavy chain 200, NF200), *ISL1* (ISLET1), *POU4F1* (BRN3A) and the mechanosensory markers *LDBH* (Lactate dehydrogenase B) and *VSN11* (Visinin Like 1) (Figure 1B). Evidence of robust DRG sensory neuronal differentiation was further supported by immunostaining analyses, showing expression of ß-III-TUBULIN and NF200 as well as a high proportion of BRN3A+ (88% and 89%) and ISLET1+ cells (85% and 89%) present within the NGN2-RUNX3 and NGN2-SHOX2 cultures, respectively (Figures 1C-1E and Table S1).

**Figure 1.**
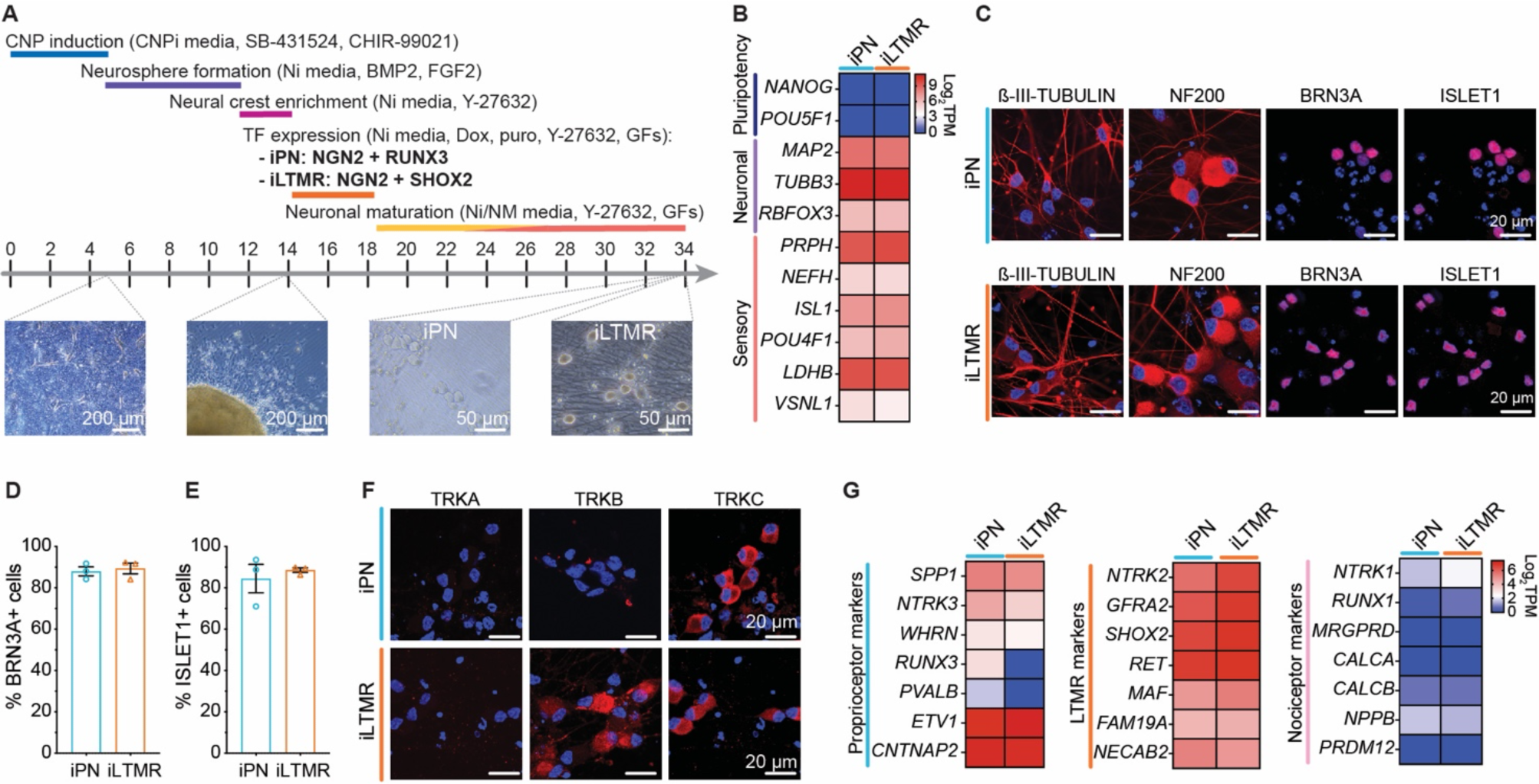
Expression of NGN2-RUNX3 or NGN2-SHOX2 in hPSC-derived neural crest progenitors induces distinct molecular profiles consistent with either proprioceptor and LTMR neurons. (A) Schematic of the protocol to derive induced-proprioceptor neurons (iPN: NGN2+RUNX3) and induced-LTMR neurons (iLTMR: NGN2+SHOX2). Growth factors (GF) include BDNF, GDNF, NT-3, and ß-NGF. (B) Heatmap of key stem cell (*NANOG*, *POU5F1*), neuron (*MAP2*, *TUBB3*, *RBFOX3*), and sensory neuron-specific (*PRPH*, *NEFH*, *ISL1*, *POU4F1*, *LDHB*, *VSNL1*) markers by bulk RNA sequencing (Log_2_TPM) of the iPN and iLTMR cultures, n = 3 biological replicates. (C) Representative immunocytochemistry images of the neuronal marker ß-III-TUBULIN (*red*) and the sensory neuronal markers NF200 (*red*), BRN3A (*red*), and ISLET1 (*red*). Nuclei are shown in blue. Scale bars = 20 μm. The percentage of iPN and iLTMR cells expressing (D) BRN3A and (E) ISLET1 compared to the number of nuclei. n = 3 biological replicates, >100 cells counted per biological replicate. (F) Representative immunocytochemistry images of the sensory neuron subtype markers TRKA (*red*), TRKB (*red*) and TRKC (*red*) in the iPN and iLTMRs. Nuclei are shown in blue. Scale bar = 20 μm. (G) Heatmap of key proprioceptor, LTMR and nociceptor markers by bulk RNA sequencing of the iPN and iLTMR cultures, presented as Log_2_TPM, n = 3 biological replicates.

To profile the transcriptional makeup of NGN2-RUNX3 and NGN2-SHOX2-induced sensory neuronal cultures, we examined gene transcripts commonly expressed in PN, LTMR and nociceptor subtypes. Reflecting PN and LTMR phenotypes, NGN2-RUNX3 neuronal cultures displayed high expression of TRKC and low levels of TRKA/TRKB, whereas the NGN2-SHOX2 neuronal cultures showed high expression of both TRKB/TRKC and low TRKA levels (Figure 1F and 1G). To determine whether our induced sensory neurons expressed transcripts associated with these mechanosensory subtypes we performed transcriptomic analyses by bulk RNA sequencing and consistently we found comparable expression levels of *ETV1*, *CNTNAP2*, *RET*, *NECAB2*, *FAM19A* and *GFRA2* between the NGN2-RUNX3 and NGN2-SHOX2 cultures (Figure 1G and Figure S2). Furthermore, there was a relatively higher expression of classic PN markers in the NGN2-RUNX3 neuronal cultures including *SPP1*, *NTRK3* (TRKC), *WHRN* (WHIRLIN), *RUNX3*, and *PVALB* (PARVALBUMIN) (Figures 1F and 1G, Figure S2). In contrast, the NGN2-SHOX2 neuronal cultures had higher expression of LTMR subtype markers such as *NTRK2* (TRKB), *GFRA2, MAF, RET*, *FAM19A*, and *NECAB2*. Additionally, minimal expression of genes associated with nociceptors was detected in both the NGN2-RUNX3 and NGN2-SHOX2-induced neurons (Figure 1G). Overall, the transcriptional profiles of the NGN2-RUNX3 and NGN2-SHOX2 cultures were consistent with previously reported PN and LTMR profiles, respectively ^11^, and thus were subsequently referred to as induced PNs (iPNs) and induced LTMRs (iLTMRs), respectively.

### iPNs and iLTMRs exhibit distinct electrophysiological signatures and firing patterns

Neuronal excitability in iPNs and iLTMRs was verified under current-clamp recording conditions, with incremental current injections (10 pA) (Figure 2, with quantification in Table S2). iPNs and iLTMRs exhibited robust action potential firing, with brief action potential duration, and increased firing frequency with increasing current stimuli (Figures 2A-2D). The induced neuron subtypes shared similar passive membrane properties, with resting membrane potentials of around −55 mV and −55 mV, capacitances of approximately 30 pF, and rheobases of about 50 pA (Figures 2E-2G). iPNs and iLTMRs fired a similar number of action potentials at two times rheobase and had an average maximum of 12 and 11 action potentials fired, respectively (Figure 2H and 2I). However, iPNs and iLTMRs displayed distinct active membrane properties, including differences in hyperpolarisation sag ratios upon hyperpolarising current injection (Figure 2J) and action potential shapes (Figures 2J-N). Specifically, at rheobase iLTMR action potentials exhibited significantly longer time to peak, rise time and action potential half-width compared to action potentials fired by iPNs (Figures 2K-2M). Moreover, iPN action potentials demonstrated faster membrane potential changes in the upstroke (larger rise slope) than action potentials fired by iLTMRs (Figure 2N). Taken together, these distinguishing features suggest differences in the molecular complement underlying the electrically activated excitability features of human-induced mechanosensory subtypes.

**Figure 2:**
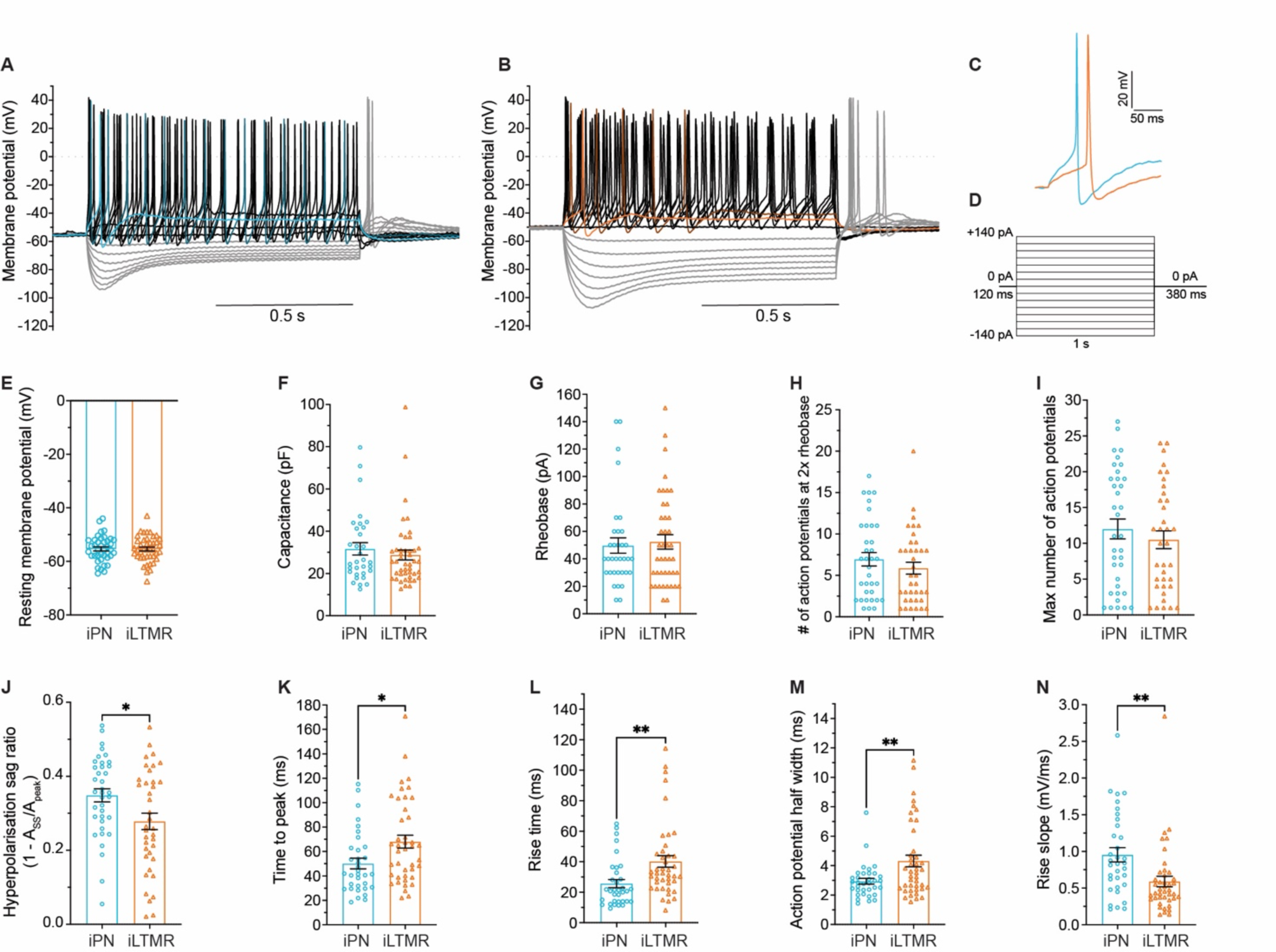
iPN and iLTMR exhibit distinct excitability profiles. Representative current-clamp recording of (A) iPN and (B) iLTMR neurons, with (C) corresponding action potentials at rheobase. (D) Membrane potential responses recorded under current-clamp conditions elicited by progressive current injections (from −140 to +140 pA, Δ 10 pA, 1 s, 0.1 Hz). Only every second trace (i.e., Λι 20 pA) is displayed for clarity. Summary of the excitability properties of the iPN and iLTMR including (E) resting membrane potential, (F) capacitance (G) rheobase, (H) the number of action potentials fired at 2× rheobase, and (I) maximum action potentials fired. (J) Hyperpolarisation sag ratio at negative current injection (−150 pA). Action potential features at rheobase include (K) time to peak, (L) rise time, (M) action potential half-width and (N) rise slope. Unpaired t-test, **p < 0.01, *p < 0.05. n = 30 – 45 neurons in total across 7 biological replicates. Data shown represent the mean ± SEM. Numeric data are included in Table S2.

The overall shape and duration of the action potential results from the complex interplay of various ion channels, including sodium, potassium, and calcium channels. Thus, we analysed voltage-gated sodium, calcium, and potassium conductance in iPNs and iLTMRs under whole-cell voltage-clamp (Figure 3, with quantification in Table S3). Voltage-gated sodium channels are crucial for mechanosensation, enabling the transmission of touch and pressure-related sensory information in neurons. iPNs and iLTMRs exhibited robust voltage-activated excitatory Na^+^ conductance (> 350 pA/pF, Figure 3B and 3C), with −36 mV and −60 mV for half-activation and half-inactivation voltages, respectively (Figure 3D and Table S3). These currents were completely inhibited by 500 nM tetrodotoxin (TTX) (Figure S3), consistent with the abundant expression of transcripts encoding TTX-sensitive (TTX-S) Na_v_ channels and low expression of transcripts (*SCN5A*, *SCN10A*, and *SCN11A*) encoding TTX-resistant (TTX-R) Na_v_ channels (Figure 3A). Furthermore, we functionally confirmed the presence of Na_v_1.1 in both induced mechanosensory subtypes, which is consistent with its expression and function in native proprioceptors and LTMRs ^4,27^ (Figures 3E-G).

**Figure 3.**
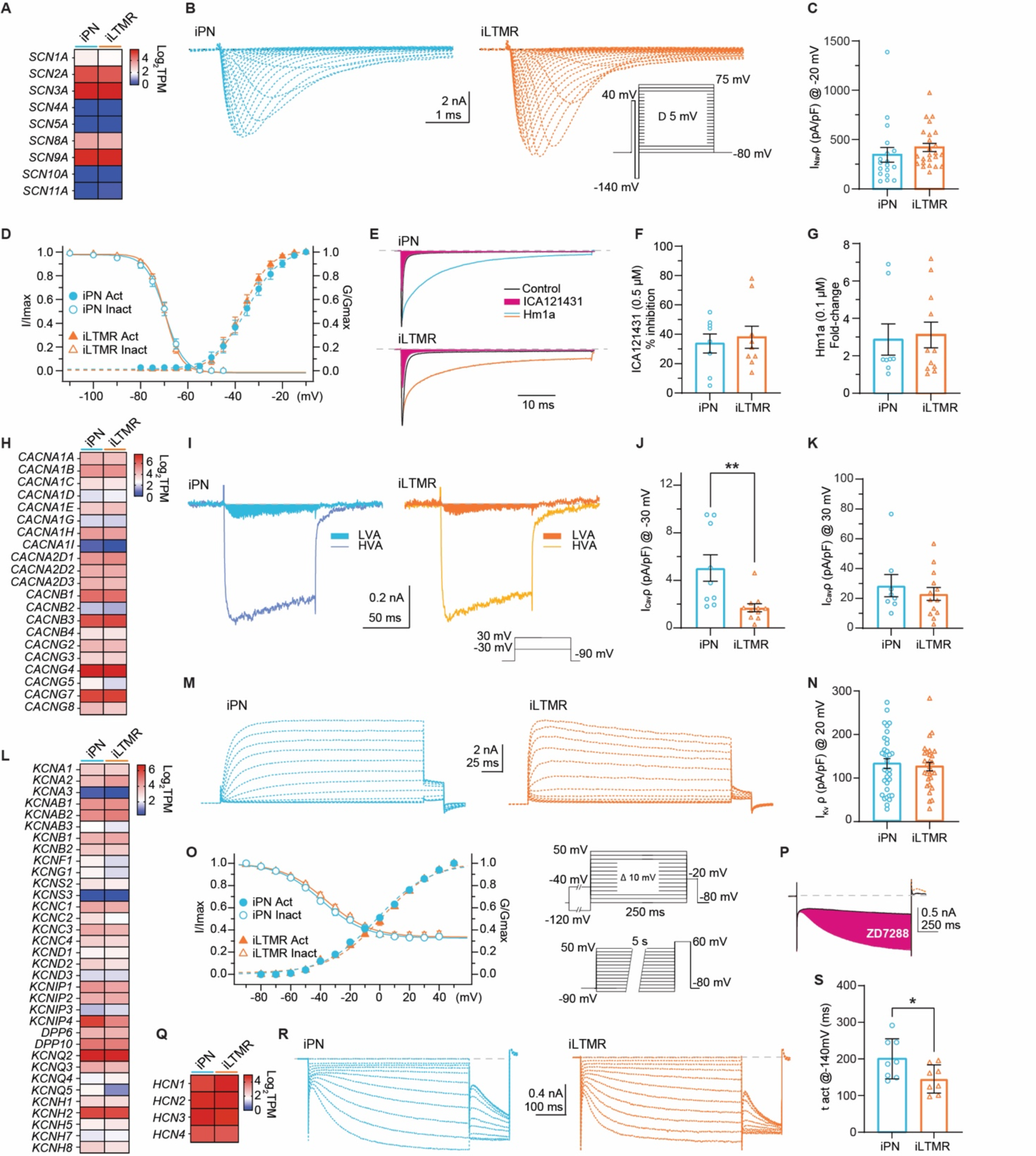
Ionic basis of excitability in iPN and iLTMR. (A-G) Depolarization-activated sodium currents (I_Nav_). (A) Transcripts encoding Na_v_ channels in iPN and iLTMR. (B) Representative Na_v_ currents in iPN and iLTMRs. (C) I_Nav_ density calculated from peak current at −20 mV. (D) Voltage-dependence activation (filled symbols) and steady-state inactivation (empty symbols) plots. (B-D) Inset shows stimulation protocols. (E) Representative examples of control (black), ICA121431 (orange), and Hm1a (blue and gold) sensitive I_Nav_ components in iPNs and iLTMRs (stimulus: 50 ms, −10 mV, Vh −80 mV, 0.1Hz). (F) Peak I_Nav_ inhibition (%) by ICA121431 (0.5 µM) in the iPNs and iLTMRs. (G) I_Nav_ enhancement by Hm1a (0.1 µM) in iPNs and iLTMRs. Accumulated charge during the stimulus in the presence of Hm1a was estimated by integrating the area under the current (AUC) trace and normalized AUC in control. (H-K) Depolarization-activated calcium currents (I_Cav_). (H) Transcripts encoding calcium channel subunits. (I-K) LVA Ca_v_-mediated currents are larger in iPNs compared to iLTMRs. (I) Representative LVA (filled) and HVA Ca_v_-mediated currents in iPN and iLTMRs. LVA (J) and HVA (K) I_Cav_ density (pA/pF) calculated from peak current (at −30 mV and 30 mV, respectively; unpaired t-test) (I-K) Inset shows stimulation protocol. (L-N) Voltage-gated potassium channels in iPNs and iLTMRs. (L) Transcripts encoding K_v_ channels and auxiliary subunits. (M) Representative I_Kv_ recorded in iPN and iLTMRs. (N) I_Kv_ density (pA/pF) calculated from peak current (at 20 mV, Vh −120 mV). (O) Voltage dependence of I_Kv_ activation (filled symbols) and steady-state inactivation (empty symbols). (M-O) Inset shows stimulation protocols. (P-S) HCN in the iPNs and iLTMRs. (P) Hyperpolarization-activated currents (I_h_) in iPNs and iLTMRs are inhibited by 10 μM ZD7288 (1 s, −140 mV, Vh −50 mV, 0.1 Hz). (Q) Transcripts encoding HCN channels in iPNs and iLTMRs. (R) I_h_ in iPNs activate slower than in iLTMRs. Families of I_h_ traces in response to 1 s hyperpolarizing steps from −50 to −140 mV (Δ10 mV; Vh −50 mV; 0.1 Hz). (S) Activation time constant (1 act) from exponential fits to I_h_ at −120 mV (unpaired t-test). Data are plotted as mean ± SEM. Numeric data are included in Table S3.

Crucial for touch and pressure sensation, voltage-gated calcium (Ca_v_) channels permit Ca^2+^ influx, influencing neurotransmitter release and facilitating signal initiation and transmission in response to mechanical stimuli. iPNs and iLTMRs exhibited abundant expression of Ca_v_ channel transcripts (Figure 3H). Notably, low voltage-activated (LVA, T-type) Ca_v_3 channels impact action potential genesis and relay triggered by low-intensity stimuli. These currents, while small, were notably larger in iPNs compared to iLTMRs during low depolarization (−30 mV) (Figure 3I, J). Based on the transcriptomes, iPN and iLTMR LVA currents correlate with the presence of transcripts of T-type channels *CACNA1G* (Ca_v_3.1), *CACNA1H* (Ca_v_3.2) and *CACNA1I* (Ca_v_3.3) (Figure 3H). Furthermore, robust high voltage-activated (HVA) Ca^2+^ currents were evoked during high depolarizations (30 mV, Vh −90 mV, Figure 3K) in both iPNs and iLTMRs (Figure 3I), aligning with the high expression of *CACNA1B* (Ca_v_2.2) and *CACNA1A* (Ca_v_2.1) transcripts (N- and P/Q-type) (Figure 3H).

Voltage-gated potassium (K_v_) channels regulate neuronal excitability and safeguard cellular homeostasis by influencing the resting membrane potential and membrane repolarization. Potassium currents are mediated by a vast family of membrane proteins actively modulating the characteristics of action potentials, including their shape, duration, and frequency. In iPNs and iLTMRs, abundant expression of multiple K_v_ channels and their modulatory subunits were detected (Figure 3L). Functionally, whole-cell patch clamp recordings of total K^+^ currents from iPN and iLTMRs revealed substantial outward currents (>125 pA/pF) largely dominated by delayed rectifier K_v_ types (Figures 3M and 3N). These currents exhibited half-activation and inactivation voltages of 0 mV and −36 mV, respectively (Figure 3O, Table S3).

Hyperpolarization-activated cyclic nucleotide-gated (HCN) channels generate hyperpolarizing currents affecting neuronal excitability and processing of mechanical stimuli. All HCN family transcripts were detected in iPNs and iLTMRs (Figure 3Q). Additionally, iPNs and iLTMRs supported robust ZD7288-sensitive (Figure 3P), hyperpolarization-activated inward currents (∼18 pA/pF), likely mediated by HCN channels, with half-activation potentials of approximately −100 mV (Figures 3R, Table S3). Notably, iPNs displayed significantly slower activation kinetics of HCN currents (at −140 mV) compared to iLTMRs under the same experimental conditions (Figure 3S), providing a plausible basis for the larger sag ratio detected in iPNs during current-clamp experiments (Figure 2J).

In summary, the absolute current densities of key voltage-gated ion channels in iPNs and iLTMRs were indistinguishable with the notable exception of apparently larger LVA Ca_v_ currents, and faster activating HCN currents in iPNs compared to iLTMRs (Figure 3 and Table S3), highlighting subtle distinguishing features between these human mechanosensory neuronal subtypes.

### The ion channel expression profiles of iPNs and iLTMRs are aligned with mechanosensory function

Bulk transcriptomic analyses of iPN and iLTMR cultures revealed abundant expression of ion channels involved in mechanosensation (Figure 4A). *PIEZO2* was highly abundant in both iPN and iLTMR cultures, which is a key mechanosensitive channel expressed in sensory neurons ^28,29^. Additionally, transcripts of channels that are associated with mechanical sensing were highly expressed in both induced neuronal subtypes, including *TMEM63B*, *TMEM120A* (TACAN), *TMEM87A* (ELKIN1), *ASIC1*, and *ASIC2* (Figure 4A). Notably, iLTMRs showed elevated *KCNK2* (TREK1) expression (Figures 4A and S3), which functions to dampen responses to mechanical stimulation ^30–32^. Conversely, iPNs exhibited higher levels of *STOML3* and *WHRN* (*Whirlin*) compared to the iLTMRs (Figure S4), which are known to regulate the sensitivity of mechanically-gated ion channels and sustained responses to stretch-evoked stimuli, respectively ^33–35^. Moreover, minimal expression of transcripts encoding ion channels associated with nociception was detected (e.g., *TRPV1*, *TRPA1*, *TRPM8*) (Figure 4A), which was further supported by negligible responses to nociceptive-like stimuli (GSK1702934A, capsaicin, menthol, AITC) compared to KCl-induced depolarization (Figures 4B and S5, with quantification in Table S4). Taken together, the iPNs and iLTMRs express a complement of ion channel transcripts relevant for sensing mechanical cues, indicative of their mechanical sensing function.

**Figure 4:**
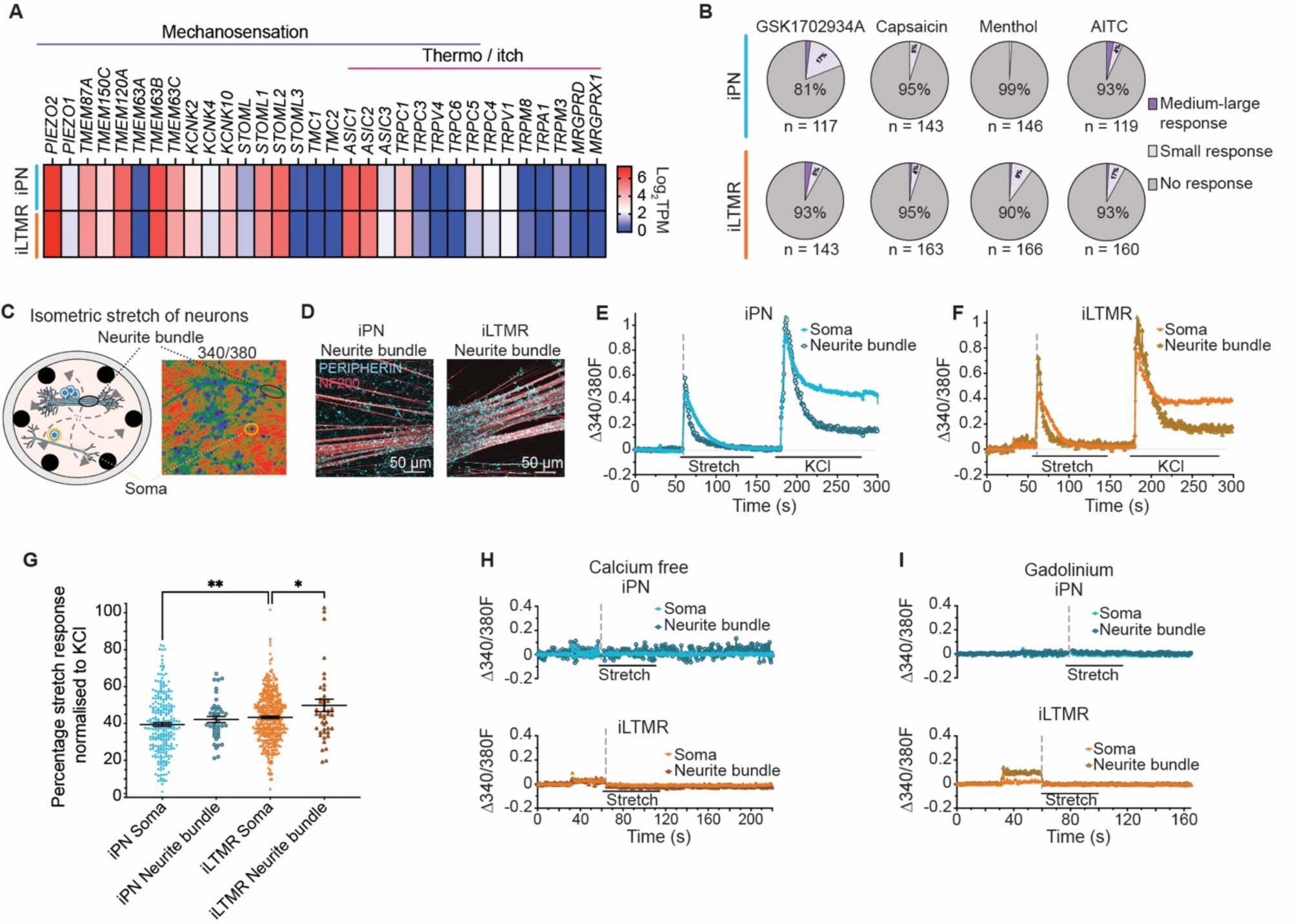
The iPN and iLTMR cultures respond to stretch-induced mechanical stimulation. (A) Heatmap of gene transcripts associated with the detection of mechanical sensations and nociception (temperature and itch) by bulk RNA sequencing (Log_2_TPM) in iPNs and iLTMRs, n = 3 biological replicates. (B) The proportion of iPNs and iLTMRs responsive to mechanical stimuli (stretch) and nociceptive stimuli (GSK1702934A, capsaicin, menthol, AITC) across n = 3 biological replicates. (C) Schematic of sensory neuron somas and neurite bundles on *IsoStretcher* PDMS chambers, measured through Fura-2 calcium imaging. (D) Representative immunocytochemistry images of PERIPHERIN (*cyan*) and NF200 (*red*) in the iPN and iLTMR neurite bundles. Scale bar = 50 μm. Representative live cell Fura-2 calcium imaging traces of an (E) iPN and (F) iLTMR soma and neurite bundle stretched by 10% using the *IsoStretcher* followed by 60 mM KCl depolarisation. Traces represent the change in the 340/380 fluorescence ratio (Δ340/380F, imaged every 0.45 s) from baseline of 1 individual neuron and 1 neurite bundle in standing CBS. (G) The percentage stretch response normalised to the KCl response. One-way ANOVA with Tukey post-hoc test, **p < 0.01, *p < 0.05, data shown represent the mean ± SEM. Representative traces of an iPN and iLTMR soma and neurite bundle incubated with (H) calcium-free CBS for 20 min before imaging and stretched by 10%. n > 80 somas and n > 10 neurite bundles across n = 3 replicates. Representative traces of an (I) iPN and iLTMR soma and neurite bundle incubated with 300 μM gadolinium for 5 min before imaging and stretched by 10%. n > 60 neurons and n > 10 neurite bundles across n = 3 replicates. Numeric data are included in Table S5.

### iPN and iLTMR respond to stretch-induced mechanical stimuli

A key role of PNs and LTMRs is their capacity to respond to multiple forms of mechanical stimulation, such as stretch and/or indentation. To explore the specific responses of iPNs and iLTMRs to different mechanical cues, we first examined their response to stretch, a low-threshold mechanical stimulus (Figure 4C). For this, we adapted the *IsoStretcher* and custom-made Polydimethylsiloxane (PDMS) chambers ^36,37^, which have been typically used to study ion channel-mediated mechanosensory transduction in cardiac cells ^38^. iPN and iLTMR neurons cultured in the PDMS chambers spontaneously clustered together to form ‘neurite bundles’, which were positive for the sensory markers PERIPHERIN and NF200 (Figures 4C and 4D). Following maturation, the neuron-laden chambers were mounted onto the *IsoStretcher* device and were isotropically stretched to 10% (chamber radial increase, ∼20% area increase). The response of both the soma and neurite bundles to stretch was recorded using Ca^2+^ imaging and quantified relative to the KCl response. Accordingly, 100% of the iPN and iLTMR somas and neurite bundles showed robust responses to the stretch-induced stimuli (Figures 4E and 4F, with quantification in Table S5). When normalised to each respective KCl response, the iPN somas and neurite bundles had comparable responses to 10% stretch whereas the iLTMR neurite bundles displayed a significantly larger response to stretch compared to iLTMR somas (Figure 4G). Interestingly, the iLTMR somas had a small but significantly larger response to stretch compared to the iPN somas (Figure 4G). Stretch-induced fluorescence changes were predominantly supported by extracellular Ca^2+^ influx, as confirmed by minimal changes in Ca^2+^-free conditions (Figure 4H and Table S5). Furthermore, inhibition of mechanosensitive channels using gadolinium ^39,40^, abolished stretch responses in both iPNs and iLTMRs (Figure 4I and Table S5). Overall, these findings support the functional mechanosensory phenotype of iPNs and iLTMRs.

### iPNs and iLTMRs exhibit distinct functional mechanosensory properties

Given the robust stretch-evoked responses of iPNs and iLTMRs, we examined whether these neurons could also respond to probe indentation to the soma as an alternative mechanical stimulus (Figure 5, with quantification in Tables S6-8). Under voltage-clamp, iPNs and iLTMRs demonstrated robust mechanically-activated (MA) whole-cell currents upon mechanical stimulation to the soma, with stimulation-intensity dependent increases in the MA current density (denoted as I_MA_) with increasing probe depth (0 – 1 μm, Δ 0.1 μm) (Figures 5A-5C). In the presence of 300 µM gadolinium, these MA currents were abolished, confirming that the response was due to activation of mechanically sensitive channels (Figure S6).

**Figure 5:**
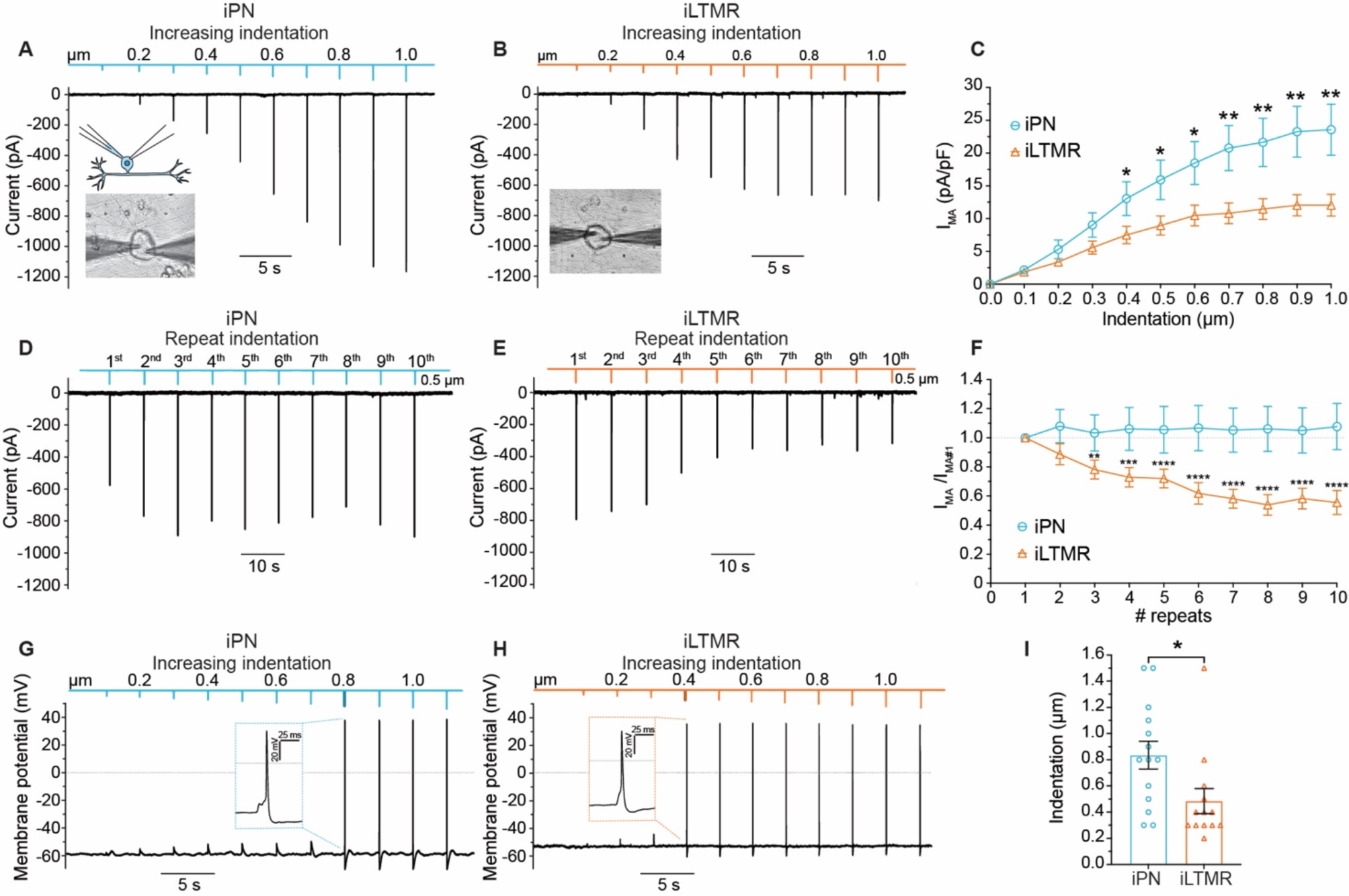
The iPN and iLTMR cultures elicit distinct stimulation-intensity dependent responses to mechanical stimulation. Representative trace of an (A) iPN and (B) iLTMR neuron under whole-cell voltage-clamp conditions with increasing 0.1 μm increments of membrane probe indentation (from 0 – 1 μm, Δ 0.1 μm, 1 s duration, 2 s rest in between indentations). Inset: iPN neuron under whole-cell patch clamp, mechanically stimulated by probe indentation. (C) The mechanically-activated (MA) current density (I_MA_) to increased mechanical stimulation by membrane probe indentation (0 – 1 μm, Δ 0.1 μm) in iPN and iLTMR. Unpaired t-test, **p < 0.01, *p < 0.05. n = 26 – 33 neurons across 5 – 8 biological replicates. Whole-cell voltage-clamp recording of a (D) iPN and (E) iLTMR neuron mechanically stimulated by repetitive 100 ms 0.5 μm membrane probe indentations followed by 5 s rest (no mechanical stimulation) 10 times (0.2 Hz). (F) The mechanically-activated current density normalised to the first probe indentation (I_MA_/I_MA#1_) following repetitive 0.5 μm indentations in iPN and iLTMRs. Unpaired t-test comparing each repeat indentation to the first indentation **p < 0.01, ***p < 0.001, ****p < 0.0001. n = 15 – 20 neurons across 4 – 5 biological replicates. Whole-cell current-clamp recording of an (G) iPN and (H) iLTMR neuron mechanically stimulated by increasing 0.1 μm increments of membrane probe indentation (from 0 – 1 μm, Δ 0.1 μm, 1 s duration, 2 s rest). Inset: first action potential firing in response to membrane probe indentation (iPN 0.8 μm, iLTMR 0.4 μm). (I) Membrane probe indentation (μm) required to elicit an action potential (mechanical rheobase). Unpaired t-test, *p < 0.05. n = 13-14 neurons across 5 biological replicates. Data shown represents the mean ± SEM. Numeric data are included in Tables S6-8.

iPNs showed heightened sensitivity to depth changes in mechanical indentations, with distinct increases in I_MA_ for each minor (0.1 μm) indentation increment over the entire range of indentations (i.e., 0.1 – 1.0 um) (Figures 5A and 5C). In contrast, the average I_MA_ for iLTMRs did not increase substantially with larger indentations of between 0.6 – 1 μm (Figure 5B and 5C). Furthermore, iPNs responded to mechanical stimulation with a substantially larger I_MA_ than iLTMRs, with approximately double that elicited in the iLTMRs at indentations > 0.4 μm (Figure 5C).

Neuronal desensitization to mechanical stimuli is crucial for regulating the responsiveness of sensory neurons to sustained stimuli. It prevents excessive activation and fatigue of mechanosensitive channels, allowing neurons to adapt to continuous mechanical input while preserving their ability to detect new stimuli. To assess desensitization, we investigated iPN and iLTMR responses to repetitive mechanical stimuli via 10 repeated probe indentations of 0.5 µm (100 ms duration) (Figure 5D and 5E). While both iPNs and iLTMRs responded to the repeated stimuli, the iPNs maintained similar response amplitudes to repeated mechanical stimulation whereas the iLTMR responses desensitised as evidenced by decreasing responses (Figures 5D and 5E). iPNs exhibited stable responses with no change in the I_MA_ amplitude with repeated mechanical stimulation compared to the initial mechanical stimulation (defined as I_MA#1_) (Figure 5F). In contrast, iLTMRs demonstrated a significant 22% decrease in the I_MA_/ I_MA#1_ after 3 repeated indentations, which continued to decrease such that after 10 repeats the response was 45% less compared to the initial mechanical stimulation (I_MA#1_) (Figure 5F).

Taken together, iPNs and iLTMRs displayed distinct responses to mechanical stimulation. iPNs elicited discrete MA and increasing responses across the whole range of mechanical stimuli, while iLTMRs’ MA currents plateaued at higher forces and desensitized with repetition.

### iPNs and iLTMRs elicit action potentials in response to mechanical stimulation with differences in their threshold of activation

A critical role of mechanosensory neurons is the ability to transduce mechanical stimuli into electrical signals, resulting in action potential firing. Thus, we sought to investigate whether iPNs and iLTMRs could recapitulate neuronal firing to mechanical stimuli by membrane probe indentation to the soma (Figures 5G and 5H). At resting membrane potential with no mechanical stimuli, iPNs and iLTMRs were silent, however, in response to 1 µm mechanical stimuli iPNs and iLTMRs elicited stereotypical single action potentials (Figure S7). Next, iPNs and iLTMRs were stimulated with progressively larger probe indentations under current-clamp (i.e., 0.1 – 1.0 um). Notably, sub-micrometer membrane indentations resulted in mechanically-elicited action potentials in both the iPN and iLTMRs somas (Figures 5G and 5H). Strikingly, iLTMRs were more excitable to mechanical stimuli with a significantly lower indentation threshold compared to iPNs, requiring an average probe indentation of 0.5 μm to elicit an action potential compared to 0.8 µm in iPNs (Figure 5I). These findings demonstrate the ability of iPNs and iLTMRs to transduce mechanical force. The differing levels of sensitivity displayed by iLTMRs and iPNs further highlight their distinguishing functions in mechanosensation.

### PIEZO2 is the major mechanically-activated sensory conductance in the human iPNs and iLTMRs

Mechanically sensitive DRG neurons can be distinguished based on the kinetics of their MA currents, which are classified as either rapidly-adapting (< 10 ms), intermediately-adapting (between 10 – 30 ms), and slowly-adapting (> 30 ms) current decays. These kinetics are determined by specific mechanosensitive channels and their modulators ^2,3,34,41^. PNs and LTMRs typically display MA currents that decay rapidly, while certain nociceptor subtypes are either mechanically insensitive or exhibit intermediately- or slowly-adapting MA current decays ^2,3,41^. We applied an exponential fit of the inactivation kinetics in iPNs and iLTMRs (Figures 6A and 6B, with quantification in Table S8) and observed that their MA currents decayed rapidly with an average time constant (τ) of 0.75 ms and 0.64 ms, respectively (Figure 6C). To compare the activation kinetics for both induced mechanosensory subtypes, the current-displacement relationship was established by normalising the I_MA_ to the observed maximal response (denoted as I_max_) and fit to a Boltzmann equation (Figure S8). The activation kinetics of iPN and iLTMR I_MA_ were similar, requiring ∼ 0.4 μm indentations to achieve half-maximal activation (I_50_) (Figure 6D). Both induced subtypes also displayed comparable mechanosensitivities with slopes of 0.16 (Figure 6E), suggesting that the same channel may be a key mediator of their mechanosensory function.

**Figure 6:**
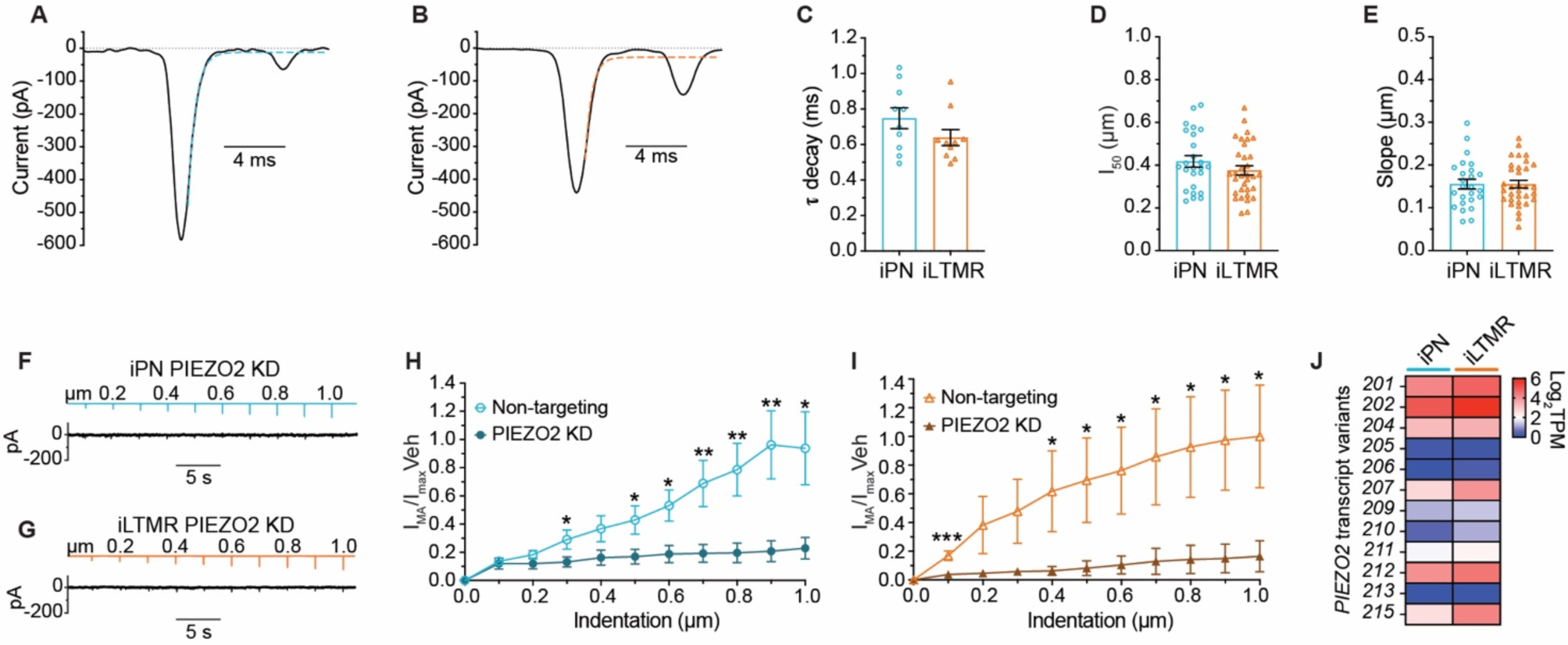
PIEZO2 is a key determinant of mechanosensory modality in iPNs and iLTMRs. Representative whole-cell voltage-clamp example of an (A) iPN and (B) iLTMR neuron mechanically stimulated by a 100 ms, 0.5 μm membrane probe indentation fit to an exponential function (dotted line). (C) Mean τ current decay of the iPNs and iLTMRs calculated from the standard exponential function of MA currents n = 10 neurons across 3 biological replicates. (D) The probe indentation to achieve 50% maximum response (I_50_) and (E) the slope of iPN and iLTMRs as calculated from the Boltzmann fit of the activation curve. n = 25 – 33 neurons across n = 5 – 8 biological replicates. Representative trace of an (F) iPN and (G) iLTMR transfected with PIEZO2 siRNA under whole-cell voltage-clamp conditions with increasing 0.1 μm increments of membrane probe indentation (from 0 – 1 μm, Δ 0.1 μm). The current density of (H) iPNs and (I) iLTMRs transfected with non-targeting siRNA and PIEZO2 siRNA in response to increased membrane probe indentation (from 0 – 1 μm, Δ 0.1 μm) normalised to the average non-targeting siRNA response at 1 μm indentation. Unpaired t-test, **p < 0.01, *p < 0.05. n = 15 – 17 neurons across 3 independent siRNA transfections. (J) Heatmap of the *PIEZO2* splice variants in iPN and iLTMR (Log_2_TPM), n = 3 biological replicates. Data shown represents the mean ± SEM. Numeric data are included in Table S9.

Given the fast MA current kinetics observed in iPNs and iLTMRs, we next examined the contribution of the PIEZO channels in mediating the iPN and iLTMR mechanosensory responses. *PIEZO2* knockdown significantly reduced iPN and iLTMR MA responses to increasing membrane displacements by up to 75% and 80%, respectively (Figures 6F-I, Table S9), highlighting the role of PIEZO2 in mediating iPN and iLTMR responses to mechanical stimulation. Furthermore, siRNA knockdown of *PIEZO1*, which is involved in the detection and transduction of mechanical itch in sensory neurons ^42^, yielded I_MA_ responses indistinguishable from non-targeting siRNA controls in both subtypes (Figure S9), indicating that PIEZO1 does not contribute to MA currents in iPNs and iLTMRs. This data suggests the potential involvement of MA channels other than PIEZO1 in iPNs and iLTMRs. Given that most of the MA current was mediated by PIEZO2, we investigated the expression of PIEZO2 splice variants between iPN and iLTMR subtypes. Notably, iPNs and iLTMRs expressed multiple *PIEZO2* splice forms, with differences in the proportions of splice variants (Figure 6J), consistent with the presence of multiple PIEZO2 splice variants within human sensory neurons^43^. Taken together, PIEZO2 is the primary mechanosensitive ion channel mediating stimulation-intensity dependent MA currents in iPNs and iLTMRs, consistent with *in vivo* PNs and LTMRs.

## Discussion

Mechanosensation is vital for touch, spatial positioning, and internal organ sensations, enabling our body to be aware and interact with our internal and external environments. Specialised mechanosensory neurons detect and discriminate a plethora of internal and external signals, however, the understanding of mechanosensory physiology has predominantly relied on rodent studies and more recently, on cadaver-explanted, axotomized DRG neurons. In this study, we generated cultures of iPN and iLTMR neurons from hPSCs and profiled their distinct molecular and functional features, which can facilitate the investigation of intrinsic cellular mechanisms within human sensory neuron mechanosensation and how this can be dysregulated in disease.

The multi-step differentiation protocol described in this study resulted in the generation of specific subpopulations of mechanosensory neurons that were highly sensitive to mechanical stimuli. A key finding is that both iPNs and iLTMRs had exclusively rapidly adapting MA current decays, mediated by PIEZO2, and did not respond to nociceptive stimuli. Furthermore, iPNs and iLTMRs had large MA current responses to < 1 µm indentations and defined responses to submicron mechanical stimulation, which has not been previously characterised in hPSC-derived neurons. Our data is in contrast to previous differentiation protocols that have induced the expression of broad sensory neuron transcription factors (e.g., NGN1, NGN2, and/or BRN3A) generating heterogeneous populations of PNs, LTMRs, and nociceptors ^16–19^. Of these, induced expression of NGN1 in hPSC-derived neural crest cells results in the generation of neurons with rapidly-adapting, intermediately-adapting, and slowly-adapting MA current decays indicative of the presence of both induced-LTMRs and induced mechanically-sensitive nociceptors ^16^. Furthermore, NGN2-alone or in combination with BRN3A in hPSC-derived neural crest cells generates mechanosensitive neurons capable of responding to nociceptive stimuli depending on the length of induced expression ^14^. Differences in the starting cell type (progenitor versus hPSC versus fibroblast), length of induced expression, and combination of small molecules, growth factors and transcription factors can dramatically alter the fate of the neurons ^12^. Importantly, our findings demonstrate the significance of inducing fate-specifying transcription factors (e.g. NGN2/RUNX3 or NGN2/SHOX2) during key developmental stages (such as neural crest) to drive neuronal differentiation to specific sensory subpopulations.

iPNs and iLTMRs revealed distinct expression and functional profiles consistent with the generation of neurons with discrete mechanosensory specializations. iPNs exhibited a remarkable sensitivity to stretch, displayed scaled responses to submicrometric membrane indentations, and sustained responses to repetitive mechanical stimuli, consistent with the high mechanical sensitivity of rodent PNs ^29,35,44–46^. Conversely, iLTMRs mirrored the characteristics of rodent rapidly adapting-LTMRs displaying desensitisation to repeated mechanical stimulation, reduced sensitivity to small changes in the depths of mechanical probe indentations, and exhibited lower mechanical thresholds for action potential firing ^44,47–50^

A limitation of our study was the inability to determine the exact subtype of PN or LTMR the cultures best fit. Typically, mammalian PNs and LTMRs are further sub-grouped based on the (1) myelination and conduction velocity, (2) location within the body, (3) end organs that the neurons innervate and associate with, (4) distinct physiological characteristics, and (5) the gene expression profile ^1,31,47^. However, within the context of our system, the comparisons of the iPN and iLTMRs to primary mechanosensory subgroups were instead based on the expression and functional characteristics of the iPNs and iLTMRs. A current challenge is to precisely determine the mechanosensory subgroups of iPNs and iLTMRs without the addition of myelination and/or innervation to end organs. Future studies to address this limitation may include conducting single-cell patch-sequencing analyses ^3,51^ and co-culturing induced neurons with end organs (e.g., muscle vs Meissner corpuscle) and glial cells, which may be necessary to further specify induced neurons into subgroups of PNs and LTMRs.

Mechanosensation is essential for everyday life, however, the exact molecular mechanisms detecting, regulating, distinguishing, and transducing different sensory stimuli between the mechanosensory neuron subtypes are still unclear. By providing an in-depth profile of human iPNs and iLTMRs, we determined that the induced neurons had differences in the excitability, responses to mechanical stimulation, and the complement of ion channels expressed. It is important to note, however, that the interaction of mechanosensory neurons with end organs is also critical for further defining and regulating responses to mechanical stimuli ^52–55^. Within the skin, LTMR axon protrusions form adherens junctions with the relevant end organs (e.g., Meissner Corpuscle vs Lanceolate ending), which serve as anchor points that, following mechanical stimulation, activate PIEZO2 allowing the neuron axons to detect changes based on the end organ they innervate ^52^. Furthermore, within duck Pacinian and Meissner corpuscles, both LTMRs and Lamellar cells respond to mechanical stimuli with distinct responses to mechanical stimulation relying on cell-cell interactions ^54,55^. The human model established in this work provides an ideal platform to investigate the interaction between induced-human mechanosensory neurons with end-organs and how they shape and fine-tune the responses of the neurons to mechanical stimulation. Intriguingly, our induced mechanosensory neurons demonstrated intrinsic differences in mechanical sensitivity and excitability even without other cell types present, such as Schwann cells, end organs, and interneurons. This highlights the significance of the molecular composition of iPNs and iLTMRs that govern their unique responses. PIEZO2 is a major channel mediating mechanosensation in iPNs and iLTMRs but its reduced expression did not convey functional differences between the two populations. Our previous studies highlighted the significant role of ELKIN1 in mechanosensation for both rodent and human DRG sensory neurons ^56^, and similar to other MA channels, expression of *ELKIN1* was comparable between iPNs and iLTMRs. The differential expression of ion channel isoforms, as observed with PIEZO2 in iPNs and iLTMRs, may underpin the functional signatures of each mechanosensory subtype. Additionally, the functional differences may be due to differences in the expression and proportions of voltage-gated ion channels ^4,31,50^, mechanosensitive channel auxiliary subunits and modulators ^33–35^, and in the lipid bilayer membrane composition and tension ^57,58^, which can enhance, modulate, or reduce the response to mechanical stimulation in mechanosensory subtypes. The differences in excitability and response to mechanical stimuli are likely due to the interplay of multiple mechanisms, which together dictate the fine-tuned differences in mechanical stimulation between human mechanosensory subgroups. Future investigations utilizing iPNs and iLTMRs in co-culture systems and/or for interrogation of candidate proteins involved in mechanosensory function will offer deeper insights into the intricate mechanisms governing human DRG sensory mechanobiology.

Loss or dysregulation of mechanosensory neuron functioning results in a range of peripheral neuropathies that can cause chronic pain, ataxia, and/or a loss of touch, bladder, stomach, and sexual sensations. By providing a comprehensive functional overview of the generated mechanosensory neurons, these cultures facilitate the screening of potential mechanosensitive modulators, ion channels, and lipid profiles. This is necessary for developing therapeutic strategies for peripheral neuropathies and to provide a greater understanding of the intrinsic cellular mechanisms within sensory neuron mechanosensation and how this can be dysregulated in peripheral neuropathies. This approach can shed light on why specific mechanosensory neurons are impacted in certain diseases whereas others remain unaffected, such as the involvement of PNs in Friedreich’s ataxia ^59,60^ and LTMRs in inflammation ^61,62^, and Autism Spectrum Disorder ^63,64^. Additionally, since PIEZO2 has a widespread role in mechanosensation it is not an ideal drug target. Alternatively, mechanosensitive channel modulators ^33,65^ may provide excellent candidates for the development of compounds that target specific mechanosensory neurons such as LTMRs but do not alter the function of the other cell types. Future work utilising activators, inhibitors, and knockdown of modulatory proteins could provide further insight into the mechanisms regulating mechanosensitivity in iPNs and iLTMRs, which in turn could identify potential targets for therapeutic intervention.

Collectively, this work describes the efficient generation and profiling of exquisitely sensitive hPSC-derived mechanosensory neurons, resembling PNs and LTMRs transcriptionally and functionally. By providing a comprehensive functional overview of iPNs and iLTMRs, this model can be used to further our understanding of human mechanosensory physiology in healthy and disease states and, in the long term, will enable the development of directed therapies toward these neuronal populations that become compromised by trauma and/or neurodegenerative conditions.

## Acknowledgements

We would like to thank Dr. Tracey Berg for her advice on plasmid design and Dr. Rachelle Balez for her advice on calcium imaging. This study was supported by funding from Friedreich’s Ataxia Research Alliance, Friedreich Ataxia Research Association, Australian Research Council Discovery Project Grants (DP240102511 and DP210102405), the University of Wollongong and an Australian Government Research Training Program (AGRTP) scholarship.

## Author contributions

Conceptualization: AJH, RKFU, MD

Methodology: AJH, JRM, NRM, SM, AT, YG, BM, RKFU, MD

Software: NRM

Validation: AJH, JRM, RKFU

Formal Analysis: AJH, JRM, NRM, AT, DK, MM, RKFU, BM, DJA, MD

Investigation: AJH, JRM, RKFU

Resources: OF, BM, DJA, MD

Data Curation: AJH, JRM, NRM, AT, RKFU

Writing – Original Draft: AJH, RKFU, MD

Writing – Review & Editing: all authors

Visualization: AJH, JRM, RKFU

Supervision: JRM, DJA, RKFU, MD

Project Administration: MD

Funding Acquisition: DJA, MD

## Declaration of interests

The authors declare no conflict of interest.

## EXPERIMENTAL MODEL AND SUBJECT DETAILS

### Lentiviral production

The lentiviral expression vectors used in this study include pLV-TetO-eGFP-PuroR (GFP control vector), pLV-TetO-hNGN2-hRUNX3-GFP-PuroR (NGN2 + RUNX3 expression vector) and pLV-TetO-hNGN2-hSHOX2-GFP-PuroR (NGN2 + SHOX2 expression vector) (Figure S1), which were designed in-house using the pLV-TetO-hNGN2-eGFP-PuroR (Addgene #79823) as a backbone. HEK293T cells were maintained at 37°C 5% CO2 in DMEM/F12 5% Foetal Bovine Serum (FBS) (SFBS – F, Interpath). For lentiviral production, HEK293T cells were passaged with Accutase (#00-4555-56, ThermoFisher) and were seeded at a density of 5,000,000 cells/T75 flask. Lentiviral particles were produced 24 h after seeding (90-100% confluence) by co-transfecting 12 µg of the plasmid encoding for the expression vector of interest, or 12 µg of the reverse tetracycline transactivator vector FUW-M2rtTA (#20342, Addgene), with the lentiviral packaging plasmids: 6 µg pMDL (#12251, Addgene), 3 µg vSVG (#8454, Addgene), and 3 µg RSV (#12253, Addgene), using polyethyleneimine (PEI) (408727, Sigma) at a ratio of 3:1 PEI:DNA in Opti-MEM (#31985062, Life technologies). The transfection media was replaced with DMEM/F12 5% FBS 6 h post transfection. Viral particles were collected at 24 h, 48 h and 72 h post-transfection and were filtered (0.45 µm pore size) and then centrifuged at 23,500 rpm for 2.5 h, at 4 °C. The pelleted viral particles were resuspended in PBS+/+ at a 200x enrichment (170 µL from 1 x T75 flask), aliquoted and stored at −80 C° until use.

### hPSC culture

All experiments were approved by the University of Wollongong Human Ethics Committee (2020/450 and 2020/451) and the University of Wollongong Institutional Biosafety Committee (GT18/03, GT19/08, GT19/09 and IBC2108). The hPSC line H9 (WA09, WiCell) was maintained on vitronectin XF™ (#07180, STEMCELL^TM^ Technologies) coated T25 flasks using TeSR-E8 media (#5990, STEMCELL^TM^ Technologies), at 37°C in a humidified incubator at 5% CO_2_. Media changes were conducted every 1 – 2 days depending on confluence and hPSCs were gently passaged, using 0.5 mM EDTA/DPBS^-^, when cultures reached a confluence of 60 – 70%.

### hPSC differentiation

hPSC differentiation to sensory neurons was based on previously published methods (Hulme 2020). Generation of sensory neurons involved the stepwise differentiation of hPSCs to caudal neural progenitors (CNPs), to neural crest spheres, to migrating neural crest cells and finally to sensory neurons, as described below (Figure 1A). Briefly, hPSCs were seeded as single cells at a density of 20,000 cells, in organ culture dishes (60 x 15mm, #353037, Corning) previously coated with 10 μg/mL laminin/PBS (#23017015, ThermoFisher) for 24 h, at 4 °C, in TeSR-E8 supplemented with 10 μM Y-27632. Following 24 h (day 1), the media was replaced with Neural Induction Media (NIM) (components outlined in Table S10), supplemented with 3 μM CHIR99021 (SML1046, Sigma), and 10 μM SB431524 (#72234, STEMCELL^TM^ Technologies). A full media change was repeated on day 3 using NIM supplemented with 3 μM CHIR99021 and 10 μM SB431524. Neurospheres were generated by harvesting day 5 CNPs. On day 5, CNPs were gently lifted using 0.5 mM EDTA/DPBS^-^ and scraping and then resuspended in Neuronal Media (NM) (components outlined in Table S10), supplemented with 20 ng/mL FGF2 (#78003, STEMCELL^TM^ Technologies) and 10 ng/mL BMP2 (RDS355BM010, In Vitro Technologies). The resuspended cell clumps were plated into ultra-low attachment U-bottom 96-well plates (100 μL/well) (CLS7007, Sigma) and the plates were centrifuged at 200 x g for 4 min. Neurosphere formation could be observed after 24 h. On day 8, 50 μL/well of NM supplemented with 20 ng/mL FGF2 and 10 ng/mL BMP2 was added, and half-media changes were conducted every 3rd day. After 7 days as neurospheres (differentiation day 12), the neurospheres were collected and plated for differentiation to sensory neurons. To enrich for migrating neural crest cells, on differentiation day 12 the neurospheres were plated as whole spheres on 12- or 13-mm glass coverslips, previously coated with 10 μg/mL Poly-D-Lysine (P6407, Sigma-Aldrich) and 10 μg/mL laminin, with NM supplemented with 10 μM Y-27632. On day 14 (48 h after neurosphere plating), the neurospheres were removed using a P200 pipette, leaving behind the migrating neural crest cells. The neural crest cells were then transduced with 1 – 2 μL/mL of lentiviral particles containing either pLV-TetO-eGFP-PuroR, pLV-TetO-hNGN2-hRUNX3-GFP-PuroR or pLV-TetO-hNGN2-hSHOX2-GFP-PuroR sequence and 1 – 2 μL/mL FUW-M2rtTA lentiviral particles for 16 h in NM supplemented with 10 μM Y-27632, 10 ng/mL BDNF (78005, STEMCELL^TM^ Technologies), 10 ng/mL GDNF (78058, STEMCELL^TM^ Technologies), 10 ng/mL NT-3 (78074, STEMCELL^TM^Technologies), and 10 ng/mL ß-NGF (78092, STEMCELL^TM^ Technologies). To remove any virus and to induce transcription factor expression, a full media change was conducted on the following day (differentiation day 15) containing 1 μg/mL doxycycline (D9891, Sigma) NM supplemented with 10 μM Y-27632, 10 ng/mL BDNF, 10 ng/mL GDNF, 10 ng/mL NT-3 and 10 ng/mL ß-NGF. Transcription factor expression was induced by the addition of doxycycline for 96 h (differentiation days 15-19). To select successfully transduced cells, 1 μg/mL puromycin (73342, STEMCELL^TM^ Technologies) was added for 48 h (day 17 – 19). To mature the progenitors into sensory neurons, media changes of NM supplemented with 10 μM Y-27632, 10 ng/mL BDNF, 10 ng/mL GDNF, 10 ng/mL NT-3 and 10 ng/mL ß-NGF were conducted every 2 – 3 days. To functionally mature the sensory neurons and mimic the nervous system’s extracellular environment BrainPhys^TM^ Neuronal Medium (BNM) (Components outlined in Table S10) was phased into the NM beginning on differentiation day 22 (25:75, 50:50, 75:25, 100:0 BNM: NM), with the same concentrations of growth factors as specified above in each media change. Between differentiation days 25 – 27, proliferating cells in the culture were removed using 2.5 μM cytosine β-D-arabinofuranoside (AraC) (C1768, Sigma) for 48 h. If neurons began to detach or cluster together, 1 μg/mL laminin was supplemented into the media to promote reattachment. The neurons were matured until day 34 and were then fixed for immunocytochemistry or harvested for total RNA extraction. Calcium imaging and patch-clamp functional analyses were performed between days 34 – 48.

## METHOD DETAILS

### Immunocytochemistry

When cultures reached the required stage for staining, the cells were washed with PBS 3 times and fixed with 4% paraformaldehyde/PBS for 20 min, at room temperature, and then PBS washed 3 times. Cells were permeabilised with 0.1% triton/PBS for 10 min and then blocked in blocking buffer (10% donkey serum/PBS (D9663, Sigma)) for 1 h at room temperature. The cultures were incubated with the appropriate primary antibody (Table S11) in blocking buffer overnight at 4°C. Following the overnight incubation, the coverslips were washed with PBS 3 times for 5 min and then incubated with the appropriate secondary antibody (Table S11) in the dark in blocking buffer for 1 h at room temperature. The secondary antibody solution was removed, and the samples were washed 3 times for 5 min in PBS and counter-stained with 1:1000 DAPI (D9542, Sigma) for 15 min. DAPI stain excess was removed after 3 repetitive 5 min PBS washes. The coverslips were mounted with a drop of ProLong™ Gold Antifade Mountant (P36934, Life Technologies Australia) onto microscope slides (MENSF41296P, ThermoFisher). Stained microscope slides were stored at 4°C in a dark microscope slide container until imaged. Images were taken using a Leica confocal SP8 microscope and exported using ImageJ (FIJI) software. Cell counts were performed using the cell count FIJI tool.

### Protein harvesting and quantification

Protein was extracted in RIPA buffer (R0278, Sigma) supplemented with protease inhibitor cocktail (P8340, Sigma). The cell lysate was centrifuged at 12,000 rpm for 20 min at 4°C and the supernatant was collected and stored at −80°C. The total protein concentration was determined via a Detergent-compatible (DC) colorimetric assay (5000112, Bio-Rad) following the manufacturer’s instructions. For Western blot analysis, the protein samples (5 µg) were resuspended in sodium dodecyl sulfate-polyacrylamide gel electrophoresis (SDS-PAGE) loading buffer (5% v/v b-mercaptoethanol (M7154, Sigma), 2× Laemmli ([0.01% v/v bromophenol blue, 25% v/v glycerol, 2% v/v SDS, 62.5 mM Tris-HCl, pH 6.8]), denatured at 95°C for 5 min and then placed on ice. Samples were separated by SDS-PAGE electrophoresis at 100 volts (V) for 1 h on 4–20% Criterion™ TGX Stain-Free™ Protein Gels (1656001, Bio-Rad) and 1x SDS-page running buffer (192 mM glycine, 3.5 mM SDS, 25 mM Tris-hydroxymethyl-methylamine). Following protein separation, the 4–20% Criterion™ TGX Stain-Free™ Protein Gel (M3148, Bio-Rad) were activated by a GelDoc XR+ (BioRad). Protein samples were transferred onto a 0.45 μm pore polyvinylidene difluoride (PVDF) membrane (IPVH00010, Millipore), previously activated in cold 100% methanol, using a Criterion blotter (1704070, Bio-Rad) at 100 V in transfer buffer (192 mM glycine, 20% v/v methanol, 25 mM tris-hydroxymethyl-methylamine) for 1.5 h. Following transfer, the membranes were washed in 0.05% Tween (P1379, Sigma) in PBS (PBST) on a rocker and imaged using GelDoc XR+ for total protein. Membranes were blocked with 10% milk/PBS rocking for 1 h, at room temperature. Membranes were incubated with the appropriate primary antibody (Table S12) in 10% milk by rocking for 16 h at 4°C. The primary antibodies were removed by washing 4 times with PBST over 20 min. The membranes were incubated with the appropriate secondary antibody (Table S12) rocking for 1 h at room temperature. The membranes were then washed 4 times with PBST and then incubated in the dark for 5 min with Clarity Western ECL substrate (1705060, Bio-Rad), and imaged using the Amersham Imager 600 (GE Industries, UK).

### RT-qPCR and bulk RNA sequencing

RNA was purified using the PureLink™ RNA Mini Kit (12183025, ThermoFisher) according to the manufacturer’s instructions. RNA concentration and quality was assessed using a Nanodrop spectrophotometer and a Bioanalyzer (Agilent). Using up to 1 µg of RNA per reaction, genomic DNA was removed, and the RNA was reverse transcribed into cDNA using the iScript™ gDNA Clear cDNA Synthesis Kit (1725035, Bio-Rad), as per the manufacturer’s guidelines. RT-qPCR was conducted using the PowerUP SYBR green master mix (A25778, ThermoFisher) in a QuantStudio 5 real-time PCR system (Applied Biosystems) using the fast run mode settings, following the manufacturer’s instructions. RNA samples that passed the quality control check (A260:280 >= 2.0, RNA and integrity number > 7.0) were utilised for sequencing. Library preparation and RNASeq analyses were performed as a service from the Garvin Institute for Medical Research (Genome One) (2× 100 base pairs, 30 million read pairs).

### Calcium imaging

When the sensory neuron cultures reached the required maturity (differentiation day 34 – 48), the neurons were incubated in Calcium imaging Bath Solution (CBS) (160 mM NaCl, 2.5 mM KCL, 5 mM CaCl_2_, 1 mM MgCl_2_, 10 mM HEPES 5 mM Glucose, pH 7.4, 320 mOsm/kg) with 6 µM Fura-2AM (F1221, ThermoFisher), and 0.04% Pluronic F-127 (P2443, Sigma) for 40 min at 37°C 5% CO_2_. Cultures were then washed with CBS, transferred to a Warner Series 20 Chamber (Warner Instruments, USA) and attached to an imaging platform (Warner Instruments, USA. Cultures were perfused with CBS at a rate of 1 mL/min using a MasterFlex C/L peristaltic pump (MasterFlex, Germany) for the duration of imaging. Calcium experiments were conducted in the dark using a DMi8 epi-fluorescent microscope (Leica Microsystems) and a dichromatic filter for dual excitation at 340 nm and 380 nm. Using the Leica calcium imaging software (LAS-X calcium imaging), the 340 and 380 channels were imaged every 0.7 s using 20x dry magnification, with imaging parameters set at 2×2 binning, 100 ms exposure, 16-bit size. Each experiment began with 2 – 4 min of CBS perfusion to establish a fluorescence baseline. The cultures were then perfused with CBS containing the required agonist followed by 60 mM KCl, with bath solution washes before and after agonist and KCl treatments. All agonists used include 100 µM Allyl isothiocyanate (AITC) (377430), 1 µM Capsaicin (M2028), 250 µM Menthol (M2772, Sigma) and 1 µM GSK1702934A (SML2323) (all from Sigma).

### Calcium imaging – stretch

hPSCs were differentiated and transduced as described in *hPSC differentiation*, however, for the isotropic stretch experiments the neural crest cells were plated onto and matured on PDMS *IsoStretcher* chambers ^36^. The stretch calcium imaging experiments were performed as described above, with modifications. Experiments were performed in static CBS bath conditions. The *IsoStretcher* ^37^ was calibrated using LabVIEW-to-Arduino interface software. Once calibrated, the chamber was fit into the *IsoStretcher* actuator. Using a Leica DMi8 microscope and the Leica calcium imaging software, the 340 and 380 channels were imaged every 0.45 s, using 20x dry magnification, with imaging parameters set at 2×2 binning, 50 ms exposure, mercury lamp setting 2, 16-bit size. To establish the baseline fluorescence intensity of the neurons and to account for the z-plane shift, the neurons in the PDMS chambers were imaged for 30s in the initial focus and then 30s in the shifted z-plane focus before stretch. Cultures on the IsoStretcher were isotropically stretched by 10% (chamber radial increase) using the LabVIEW-to-Arduino interface software. A final maximal depolarization was achieved by KCl addition (final concentration of 60 mM) to the PDMS chamber 3 min post-stretch. For the gadolinium experiments, chambers were incubated in 300 µM gadolinium (Gadolinium (III) chloride hexahydrate, 13450-84-5, Sigma) for 5 min before imaging and stretching. For the extracellular calcium-free control, the CBS solution was exchanged with a calcium-free CBS (10 mM EGTA, 162 mM NaCl, 2.5 mM KCL, 1 mM MgCl_2_, 10 mM HEPES 5 mM Glucose, pH 7.4, 320 mOsm/kg) and imaging experiments proceeded as described above.

### Electrophysiology

Whole-cell patch-clamp recordings were performed at room temperature (20 – 24°C) using an inverted microscope (Nikon) and a MultiClamp 700B Amplifier, digitized with a Digidata 1440 and controlled with pClamp11 software (Molecular Devices, San Jose, CA, USA). The bath solution for current clamp experiments was made with 135 mM NaCl, 2 mM CaCl_2_, 2 mM MgCl_2_, 5 mM KCl, 10 mM glucose, 10 mM HEPES, pH 7.4, osmolality 315± 5 mOsm/kg. For voltage-clamp experiments, the bath solution varied depending on the ion channel being examined. For K^+^ currents (I_K_) the bath solution above was supplemented with 1 μM TTX (Tetrodotoxin Citrate, Abcam (Melbourne, Australia)). Na^+^ currents (I_Na_) were isolated using a bath solution containing 110 mM NaCl, 2 mM CaCl_2_, 2 mM MgCl_2_, 30 mM TEA-Cl, 10 mM D-Glucose, and 10 mM HEPES, pH 7.3. To isolate Ca^2+^ currents (I_Ca_), the extracellular solution contained 140 mM TEA-Cl, 10 mM CaCl_2_, 1 mM MgCl_2_, 10 mM HEPES, 10 mM D-Glucose, pH 7.3. Borosilicate glass patch pipettes (World Precision Instruments, USA) were pulled using a P-97 Flaming/Brown micropipette puller (Sutter Instruments), fire polished to resistance between 2 – 4 MΩ and filled with intracellular buffer (140 mM K-gluconate, 10 mM NaCl, 2 mM MgCl_2_, 10 mM HEPES, 5 mM EGTA, pH 7.2, osmolality 295± 5 mOsm/kg). Whole-cell recording configuration was obtained under voltage-clamp settings. Series resistance was compensated for at ≥ 60%, and whole-cell currents were sampled at 100 kHz and filtered to 10 kHz. Neuronal excitability was assessed under current-clamp conditions. Action potential firing was elicited by 1 s incremental (10 pA) current injections (−150 pA to 150 pA). Ion channel modulators (Gadolinium, ICA 121431, Hm1a, ZD7288) were applied to cells using whole bath perfusion.

Mechanical stimulation of the neurons was achieved by probe indention using a fire-sealed borosilicate glass patch pipette (denoted as probe) placed at an angle of 45° (to the supporting glass coverslip), filled with intracellular buffer connected to a micromanipulator (PatchStar) controlled by LinLab 2 software (Scientifica). The standard mechanically-activated (MA) stimulation protocol consisted of 10 incremental 0.1 µm, 1 s long indentations delivered up to a maximal 1 µm indentation every 2 s (0.1 µm – 1.0 µm, Δ 0.1 µm, 0.5 Hz), recorded in gap-free voltage-clamp or current-clamp conditions, variations in the stimulation protocol are described in the respective figure legends.

### siRNA transfection or Knockdown of PIEZO1 and PIEZO2

Neurons were transfected using Accell SMARTpool siRNA (knockdown day 1) with either 1 µM non-targeting siRNA (D-001960-01-20, Accell, Horizon Discovery Group Company), 1 µM human SMARTpool PIEZO1 siRNA (E-020870-00-050, Accell, Horizon Discovery Group Company) or 1 µM PIEZO2 (E-013925-00-0050, Accell, Horizon Discovery Group Company) in BrainPhys Media supplemented with 10 μM Y-27632, 10 ng/mL BDNF, 10 ng/mL GDNF, 10 ng/mL NT-3 and 10 ng/mL ß-NGF. Following 48 h transfection (knockdown day 3), a full media change containing fresh siRNAs was conducted. The neurons were whole-cell patch clamped 72 h post initial siRNA transfection (KD day 4). To confirm siRNA knockdown, neurons were harvested for RNA, and RT-qPCR was conducted. To determine the mechanical response following transfection, the neurons were whole-cell voltage clamped using borosilicate glass patch pipettes fire polished to a resistance between 2 – 4 MΩ and filled with CsF-intracellular buffer (110 mM CsF, 30 mM CsCl, 10 mM NaCl, 2 mM MgCl_2_, 10 mM HEPES and 5 mM EGTA, pH 7.2, osmolarity 295± 5 mOsm/kg).

## QUANTIFICATION AND STATISTICAL ANALYSIS

### Bulk RNA sequencing analysis

The initial RNAseq processing involved the utilization of DRAGEN RNA Pipeline 3.7.5. Following adapter trimming, RNAseq reads with a Phred Quality score greater than 20 were retained and preprocessed. The input reads ranged from 124 million to 190 million, with paired proportions falling within the range of 94.7% to 96.3% and median insert sizes ranging from 115 to 137. During the parameter selection phase, the GRCh38 reference was chosen, excluding alternative contigs and including decoy, while the gencode_grch38.v32.annotation.gtf was employed for RNA annotation purposes. The DRAGEN RNA pipeline utilized the DRAGEN RNA-Seq spliced aligner. After obtaining the raw count matrix, rows with zero expression were removed. ComBat_seq ^66^ normalization was applied to correct potential biases due to batches of sub-cultures. Further, Gene differential expression analysis was performed using Deseq2 ^67^. The average data from 3 biological replicates was presented as the log2 Transcripts per Million (Log2TPM).

### Calcium imaging analysis

Calcium imaging data was analysed using LAS-X calcium imaging software. Traces were generated by calculating the change in the 340/380 fluorescence ratio subtracted by the baseline fluorescence of each cell. For the stretch experiments, due to some soma and neurites shifting in and out of the field of view once the stretch occurred, the baseline fluorescence of each cell was taken as the average fluorescence intensity of 20 s before KCl addition. The maximum of the responses to the agonist and KCl were calculated using a custom script written on Python software available at: https://doi.org/10.5281/zenodo.7460899. The percentage response was calculated by:

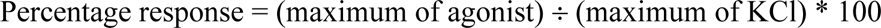

To determine the proportion of responsive cells, a signal-to-noise cut-off threshold of 2% (of maximal response) was set as values below the threshold were indistinguishable from noise and thus classified as “no response”.

### Electrophysiology analysis

All traces and action potentials were analysed and exported using Clampfit version 11.1.0.23. The rheobase was defined as the minimum amount of current necessary to evoke a single action potential. Action potential shape analysis was based on the rheobase action potential. MA currents were pre-processed using in-built Clampfit functions by (1) baselining to the average current level recorded 30 ms before stimulation and (2) filtering using a Low-pass Gaussian filter (Cut-off 770 Hz, 13 coefficients). Mechanoclamp recordings were exported into comma separated value files and the maximum peak was calculated using a custom-developed Python script available at https://doi.org/10.5281/zenodo.7460899. To calculate the mean τ current decay, MA currents were recorded in gap-free voltage-clamp conditions with a 0.5 µm (100 ms duration) membrane probe indentation and the MA current was fit to a standard exponential function:

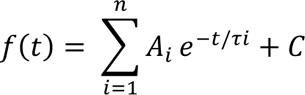

To compare the mechanical activation kinetics of the neurons, the I_MA_ was normalised to the maximum response (denoted as I_MAX_) for each neuron (denoted as I_MA_/I_MAmax_) and was fit to a Boltzmann sigmoidal with the constraints ‘top = 1, bottom = 0’ generating a ‘mechanical dependence of activation’ curve. Using the mechanical dependence of activation curve, the probe indentation required to achieve 50% of the maximum response (Indent_50_) and the mechanosensitivity (slope) was determined.

Voltage dependence of activation and steady-state inactivation (SSI) of the various ionic currents were fit by the modified Boltzmann equation:

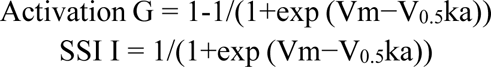

where I is the current, G is the conductance, Vm is the pre-pulse potential, V_0.5_ is the half-maximal activation potential and ka is the slope factor.

### Statistical analysis

All statistical analysis was performed and presented using GraphPad Prism 9 unless stated otherwise. Data was presented as the mean ± standard error of the mean (SEM). Groups were compared using either Student’s T-test or One-Way Analysis of Variance (ANOVA) with Tukey post-hoc test to determine statistical significance. The specific statistical method and number of replicates are specified in the relevant figure legends and tables.

## Supplementary tables

**Table S1:** Quantification of the percentage of BRN3A+ and ISLET1+ cells in each biological replicate.

|  | iPN |  |  | iLTMR |  |  |  |
| --- | --- | --- | --- | --- | --- | --- | --- |
|  | Mean | SEM | n | Mean | SEM | n | Unpaired t-test |
| BRN3A+ cells (%) | 88 | 2.175 | 3 | 89.37 | 2.628 | 3 | 0.7088 |
| ISLET1+ cells (%) | 84.48 | 6.853 | 3 | 88.59 | 1.11 | 3 | 0.5856 |

^n = biological replicates

**Table S2:** Quantification of the iPN and iLTMR excitability profiles.

|  | iPN |  |  | iLTMR |  |  |  |
| --- | --- | --- | --- | --- | --- | --- | --- |
|  | Mean | SEM | n | Mean | SEM | n | Unpaired t-test |
| Resting membrane potential (mV) | -55.4 | 0.757 | 40 | -55.4 | 0.746 | 42 | 0.9995 |
| Capacitance (pF) | 31.64 | 2.91 | 32 | 28.75 | 2.373 | 45 | 0.4411 |
| Rheobase (pA) | 49.71 | 5.609 | 34 | 52.38 | 5.222 | 42 | 0.729 |
| Number of action potentials at 2 times rheobase | 6.943 | 0.8106 | 35 | 5.865 | 0.7209 | 37 | 0.3226 |
| Maximum action potentials fired | 12 | 1.39 | 36 | 10.5 | 1.24 | 36 | 0.4223 |
| Hyperpolarisation sag ratio at -140pA | 0.3484 | 0.01779 | 37 | 0.2782 | 0.0223 | 38 | **0.0166** |
| Time to peak (ms) | 50.15 | 4.325 | 34 | 68.05 | 5.32 | 42 | **0.0136** |
| Rise time (ms) | 25.66 | 2.654 | 34 | 40.21 | 3.824 | 42 | **0.0039** |
| Action potential half-width (ms) | 2.94 | 0.1985 | 34 | 4.32 | 0.3896 | 42 | **0.0043** |
| Rise slope (mV/ms) | 0.9534 | 0.0964 | 34 | 0.5887 | 0.0715 | 42 | **0.0027** |
| Peak amplitude (mV) | 94.79 | 1.875 | 34 | 95.11 | 1.747 | 42 | 0.9016 |

^n = neurons

**Table S3:** Quantification of the ionic basis of excitability in iPN and iLTMR.

|  | iPN |  |  | iLTMR |  |  |  |
| --- | --- | --- | --- | --- | --- | --- | --- |
|  | Mean | SEM | n | Mean | SEM | n | Unpaired t-test |
| $I_{NaV}$ (pA/pF) | -345.5 | 74.6 | 18 | -420.5 | 41.8 | 24 | 0.3572 |
| $Na_v$ act $V_{0.5}$ (mV) | -35.6 | 1.7 | 18 | -37.1 | 1.1 | 25 | 0.4352 |
| $Na_v$ inact $V_{0.5}$ (mV) | -61.9 | 1.9 | 17 | -60.0 | 1.1 | 17 | 0.4116 |
| $Na_v$ TTX block (%) | 99.4 | 0.25 | 6 | 99.4 | 0.1 | 6 | 1 |
| $Na_v$ ICA block (%) | 33.8 | 6.5 | 8 | 38.0 | 7.5 | 9 | 0.6789 |
| $Na_v$ Hm1a fold change | 2.9 | 0.8 | 8 | 3.1 | 0.7 | 11 | 0.8224 |
| HVA $I_{Cav}$ (pA/pF) | -37.5 | 11.1 | 9 | -23.0 | 4.4 | 13 | 0.5026 |
| LVA $I_{Cav}$ (pA/pF) | -5.0 | 1.1 | 9 | -3.2 | 1.3 | 13 | **0.0058** |
| $I_{Kv}$ (pA/pF) | 128.2 | 11.6 | 35 | 125.2 | 9.8 | 29 | 0.8467 |
| $K_v$ act $V_{0.5}$ (mV) | 3.8 | 2.3 | 22 | 5.0 | 1.9 | 32 | 0.7131 |
| $K_v$ inact $V_{0.5}$ (mV) | -37.3 | 1.9 | 34 | -35.9 | 1.9 | 29 | 0.6113 |
| $I_h$ (pA/pF) | -19.0 | 3.5 | 11 | -17.1 | 2.7 | 10 | 0.6703 |
| $I_h$ act $V_{0.5}$ (mV) | -98.4 | 2 | 11 | -101.6 | 1.8 | 9 | 0.2758 |
| $I_h$ tau act (ms) | 145.3 | 13.6 | 8 | 200.5 | 19.3 | 8 | **0.0351** |

^n = neurons

**Table S4:**
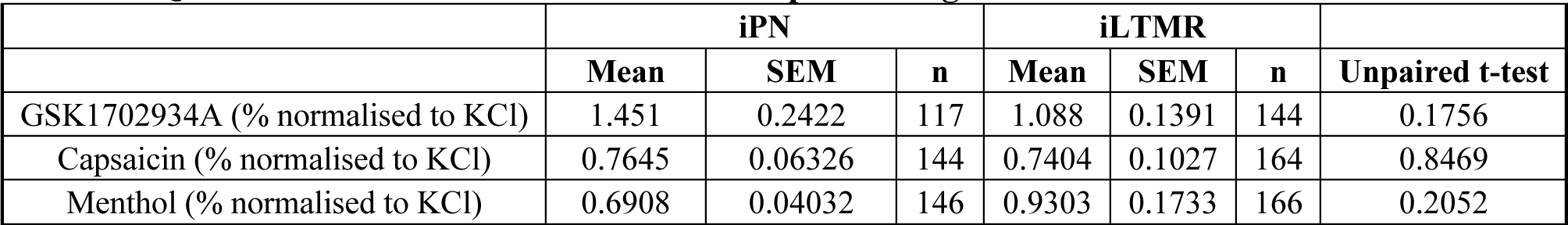
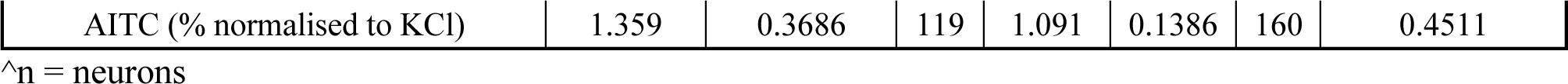
Quantification of the iPN and iLTMR response to agonists.

|  | iPN |  |  | iLTMR |  |  |  |
| --- | --- | --- | --- | --- | --- | --- | --- |
|  | Mean | SEM | n | Mean | SEM | n | Unpaired t-test |
| GSK1702934A (% normalised to KCl) | 1.451 | 0.2422 | 117 | 1.088 | 0.1391 | 144 | 0.1756 |
| Capsaicin (% normalised to KCl) | 0.7645 | 0.06326 | 144 | 0.7404 | 0.1027 | 164 | 0.8469 |
| Menthol (% normalised to KCl) | 0.6908 | 0.04032 | 146 | 0.9303 | 0.1733 | 166 | 0.2052 |
| AITC (% normalised to KCl) | 1.359 | 0.3686 | 119 | 1.091 | 0.1386 | 160 | 0.4511 |

^n = neurons

**Table S5:**
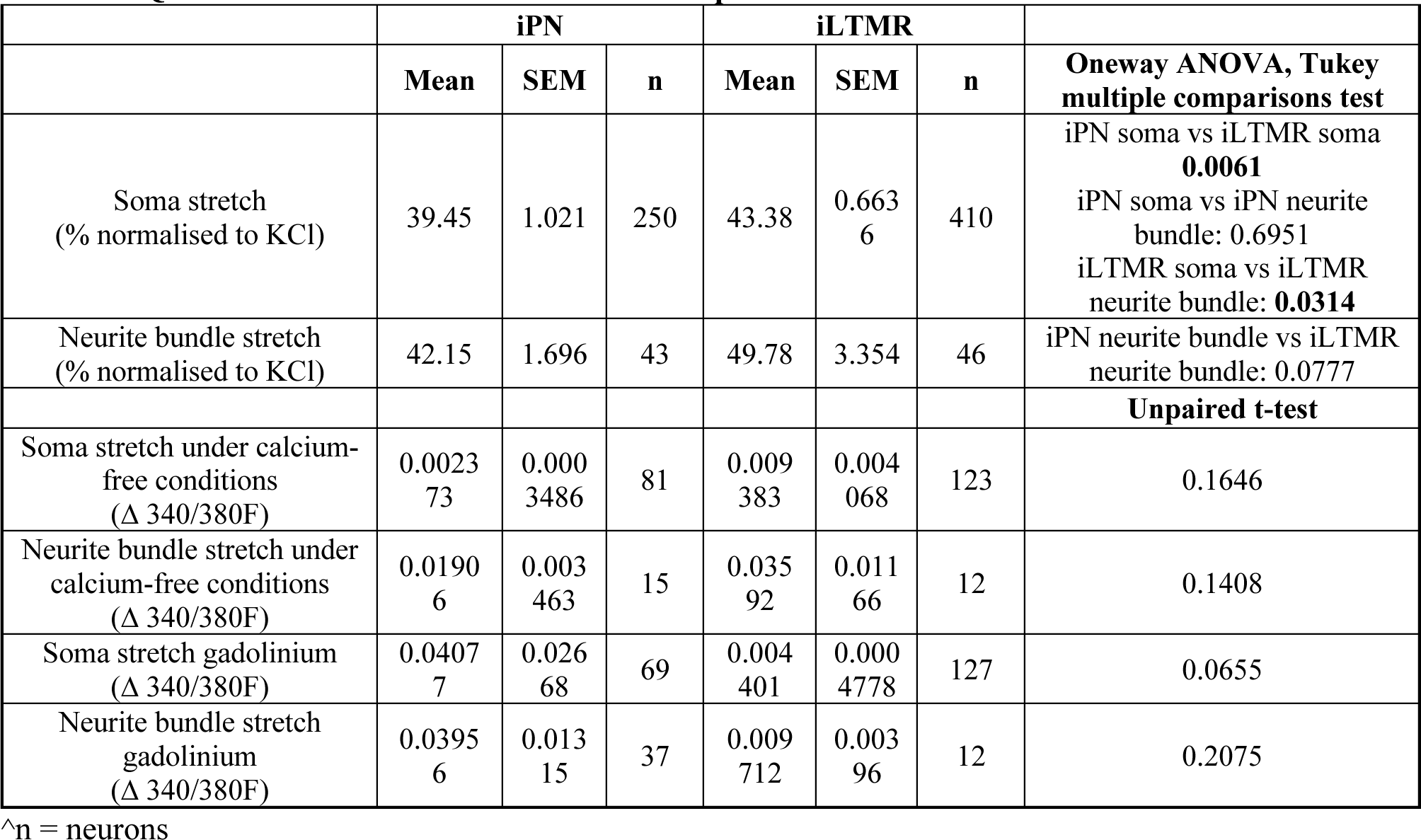
Quantification of the iPN and iLTMR response to stretch.

**Table S6:** Quantification of the response of iPN and iLTMR to increasing probe indentation.

|  | iPN |  |  | iTMR |  |  |  |
| --- | --- | --- | --- | --- | --- | --- | --- |
|  | Mean | SEM | n | Mean | SEM | n | Unpaired t-test |
| I<sub>MA</sub> at 0.1 μm (pA/pF) | 2.169 | 0.4408 | 26 | 1.85 | 0.2669 | 33 | 0.5199 |
| I<sub>MA</sub> at 0.2 μm (pA/pF) | 5.312 | 1.426 | 26 | 3.382 | 0.5582 | 33 | 0.1769 |
| I<sub>MA</sub> at 0.3 μm (pA/pF) | 9.028 | 1.867 | 26 | 5.582 | 0.9993 | 33 | 0.0905 |
| I<sub>MA</sub> at 0.4 μm (pA/pF) | 13.05 | 2.576 | 26 | 7.501 | 1.318 | 33 | **0.0464** |
| I<sub>MA</sub> at 0.5 μm (pA/pF) | 15.92 | 3.003 | 26 | 8.946 | 1.449 | 33 | **0.0295** |
| I<sub>MA</sub> at 0.6 μm (pA/pF) | 18.47 | 3.272 | 26 | 10.47 | 1.578 | 33 | **0.0222** |
| I<sub>MA</sub> at 0.7 μm (pA/pF) | 20.75 | 3.449 | 26 | 10.81 | 1.582 | 33 | **0.0068** |
| I<sub>MA</sub> at 0.8 μm (pA/pF) | 21.63 | 3.684 | 26 | 11.45 | 1.584 | 33 | **0.0077** |
| I<sub>MA</sub> at 0.9 μm (pA/pF) | 23.25 | 3.866 | 25 | 12.04 | 1.62 | 33 | **0.005** |
| I<sub>MA</sub> at 1.0 μm (pA/pF) | 23.57 | 3.891 | 25 | 12.03 | 1.663 | 32 | **0.0047** |

^n = neurons

**Table S7:** Quantification of the response of iPN and iLTMR to repeated probe indentation.

|  | iPN |  |  |  | iTMR |  |  |  |
| --- | --- | --- | --- | --- | --- | --- | --- | --- |
|  | Mean | SEM | n | Unpaired t-test | Mean | SEM | n | Unpaired t-test |
| I<sub>MA</sub> /I<sub>max#1</sub> at 2nd repeat | 1.08 | 0.1145 | 15 | 0.4908 | 0.8867 | 0.07254 | 20 | 0.1265 |
| I<sub>MA</sub> /I<sub>max#1</sub> at 3rd repeat | 1.032 | 0.1241 | 15 | 0.7954 | 0.7817 | 0.0651 | 20 | **0.0018** |
| I<sub>MA</sub> /I<sub>max#1</sub> at 4th repeat | 1.061 | 0.1468 | 15 | 0.68 | 0.7284 | 0.06629 | 20 | **0.0002** |
| I<sub>MA</sub> /I<sub>max#1</sub> at 5th repeat | 1.056 | 0.1604 | 15 | 0.7312 | 0.7199 | 0.06402 | 20 | **<0.0001** |
| I<sub>MA</sub> /I<sub>max#1</sub> at 6th repeat | 1.067 | 0.1556 | 15 | 0.6703 | 0.618 | 0.07315 | 20 | **<0.0001** |
| I<sub>MA</sub> /I<sub>max#1</sub> at 7th repeat | 1.053 | 0.1517 | 15 | 0.7311 | 0.581 | 0.06438 | 20 | **<0.0001** |
| I<sub>MA</sub> /I<sub>max#1</sub> at 8th repeat | 1.061 | 0.1548 | 14 | 0.6851 | 0.5386 | 0.07102 | 20 | **<0.0001** |
| I<sub>MA</sub> /I<sub>max#1</sub> at 9th repeat | 1.05 | 0.1553 | 14 | 0.7401 | 0.5819 | 0.07092 | 20 | **<0.0001** |
| $I_{MA}/I_{max\#1}$ at 10th repeat | 1.08 | 0.16 | 12 | 0.5929 | 0.5549 | 0.08256 | 19 | <0.0001 |

^n = neurons

**Table S8:** Quantification of the response of iPN and iLTMR to probe indentation.

|  | iPN |  |  | iLTMR |  |  | Unpaired t-test |
| --- | --- | --- | --- | --- | --- | --- | --- |
|  | Mean | SEM | n | Mean | SEM | n |  |
| Indentation required to elicit an action potential ($\mu\text{m}$) | 0.84 | 0.11 | 14 | 0.48 | 0.095 | 13 | **0.0218** |
| t current decay (ms) | 0.75 | 0.059 | 10 | 0.64 | 0.045 | 10 | 0.16 |
| $I_{50}$ ($\mu\text{m}$) | 0.42 | 0.027 | 25 | 0.38 | 0.022 | 33 | 0.221 |
| Slope ($\mu\text{m}$) | 0.16 | 0.011 | 25 | 0.16 | 0.0088 | 33 | 0.98 |

^n = neurons

**Table S9:**
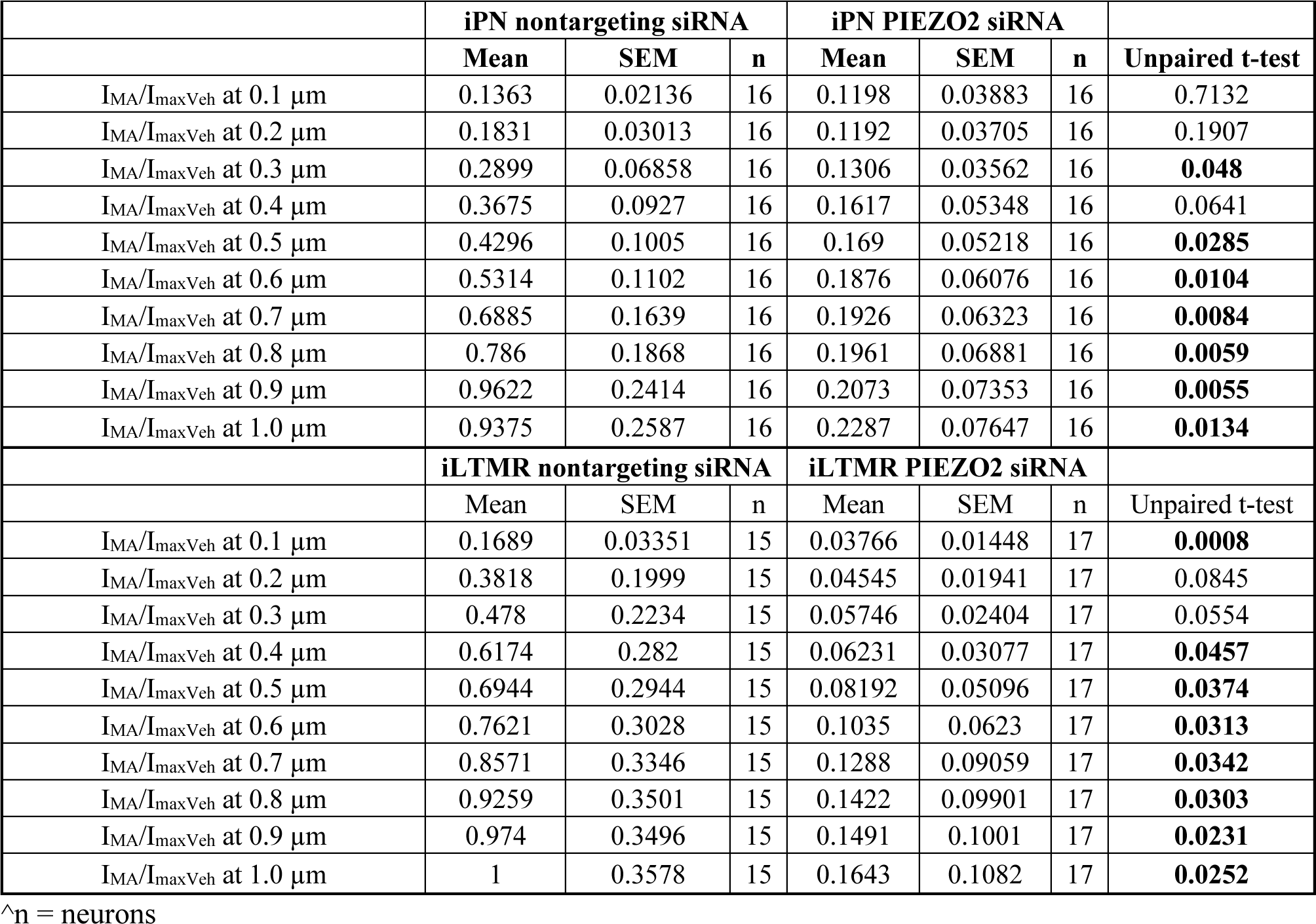
Quantification of the response of iPN and iLTMR to increasing probe indentation following PIEZO2 KD.

|  | iPN nontargeting siRNA |  |  | iPN PIEZO2 siRNA |  |  | Unpaired t-test |
| --- | --- | --- | --- | --- | --- | --- | --- |
|  | Mean | SEM | n | Mean | SEM | n |  |
| $I_{MA}/I_{maxVeh}$ at 0.1 $\mu\text{m}$ | 0.1363 | 0.02136 | 16 | 0.1198 | 0.03883 | 16 | 0.7132 |
| $I_{MA}/I_{maxVeh}$ at 0.2 $\mu\text{m}$ | 0.1831 | 0.03013 | 16 | 0.1192 | 0.03705 | 16 | 0.1907 |
| $I_{MA}/I_{maxVeh}$ at 0.3 $\mu\text{m}$ | 0.2899 | 0.06858 | 16 | 0.1306 | 0.03562 | 16 | **0.048** |
| $I_{MA}/I_{maxVeh}$ at 0.4 $\mu\text{m}$ | 0.3675 | 0.0927 | 16 | 0.1617 | 0.05348 | 16 | 0.0641 |
| $I_{MA}/I_{maxVeh}$ at 0.5 $\mu\text{m}$ | 0.4296 | 0.1005 | 16 | 0.169 | 0.05218 | 16 | **0.0285** |
| $I_{MA}/I_{maxVeh}$ at 0.6 $\mu\text{m}$ | 0.5314 | 0.1102 | 16 | 0.1876 | 0.06076 | 16 | **0.0104** |
| $I_{MA}/I_{maxVeh}$ at 0.7 $\mu\text{m}$ | 0.6885 | 0.1639 | 16 | 0.1926 | 0.06323 | 16 | **0.0084** |
| $I_{MA}/I_{maxVeh}$ at 0.8 $\mu\text{m}$ | 0.786 | 0.1868 | 16 | 0.1961 | 0.06881 | 16 | **0.0059** |
| $I_{MA}/I_{maxVeh}$ at 0.9 $\mu\text{m}$ | 0.9622 | 0.2414 | 16 | 0.2073 | 0.07353 | 16 | **0.0055** |
| $I_{MA}/I_{maxVeh}$ at 1.0 $\mu\text{m}$ | 0.9375 | 0.2587 | 16 | 0.2287 | 0.07647 | 16 | **0.0134** |
|  | iLTMR nontargeting siRNA |  |  | iLTMR PIEZO2 siRNA |  |  | Unpaired t-test |
|  | Mean | SEM | n | Mean | SEM | n |  |
| $I_{MA}/I_{maxVeh}$ at 0.1 $\mu\text{m}$ | 0.1689 | 0.03351 | 15 | 0.03766 | 0.01448 | 17 | **0.0008** |
| $I_{MA}/I_{maxVeh}$ at 0.2 $\mu\text{m}$ | 0.3818 | 0.1999 | 15 | 0.04545 | 0.01941 | 17 | 0.0845 |
| $I_{MA}/I_{maxVeh}$ at 0.3 $\mu\text{m}$ | 0.478 | 0.2234 | 15 | 0.05746 | 0.02404 | 17 | 0.0554 |
| $I_{MA}/I_{maxVeh}$ at 0.4 $\mu\text{m}$ | 0.6174 | 0.282 | 15 | 0.06231 | 0.03077 | 17 | **0.0457** |
| $I_{MA}/I_{maxVeh}$ at 0.5 $\mu\text{m}$ | 0.6944 | 0.2944 | 15 | 0.08192 | 0.05096 | 17 | **0.0374** |
| $I_{MA}/I_{maxVeh}$ at 0.6 $\mu\text{m}$ | 0.7621 | 0.3028 | 15 | 0.1035 | 0.0623 | 17 | **0.0313** |
| $I_{MA}/I_{maxVeh}$ at 0.7 $\mu\text{m}$ | 0.8571 | 0.3346 | 15 | 0.1288 | 0.09059 | 17 | **0.0342** |
| $I_{MA}/I_{maxVeh}$ at 0.8 $\mu\text{m}$ | 0.9259 | 0.3501 | 15 | 0.1422 | 0.09901 | 17 | **0.0303** |
| $I_{MA}/I_{maxVeh}$ at 0.9 $\mu\text{m}$ | 0.974 | 0.3496 | 15 | 0.1491 | 0.1001 | 17 | **0.0231** |
| $I_{MA}/I_{maxVeh}$ at 1.0 $\mu\text{m}$ | 1 | 0.3578 | 15 | 0.1643 | 0.1082 | 17 | **0.0252** |

^n = neurons

**Table S10.** Media and Components.

| Neural Induction Media |  |  |
| --- | --- | --- |
| Reagent | Catalogue number | Company |
| Neurobasal medium | 21103-049 | ThermoFisher |
| DMEM/F12 |  | Made in house |
| N-2 supplement (100x) | 17502-048 | ThermoFisher |
| B-27 supplement without vitamin A (50x) | 12587-010 | ThermoFisher |
| Insulin-transferrin-Selenium-A (100x) | 51300-044 | ThermoFisher |
| 2 mM L-glutamine | 25030149 | Life Technologies |
| 0.3% glucose | G8769 | Sigma |
| **Neuronal Media** |  |  |
| **Reagent** | **Catalogue number** | **Company** |
| Neurobasal medium | 21103-049 | ThermoFisher |
| N-2 supplement (100x) | 17502-048 | ThermoFisher |
| B-27 supplement without vitamin A (50x) | 12587-010 | ThermoFisher |
| Insulin-transferrin-Selenium-A (100x) | 51300-044 | ThermoFisher |
| L-glutamine (100x) | 25030149 | Life Technologies |
| **BrainPhys media** |  |  |
| **Reagent** | **Catalogue number** | **Company** |
| BrainPhys™ Neuronal Medium | 5790 | STEMCELL™ Technologies |
| NeuroCult™ SM1 Without Vitamin A | 5731 | STEMCELL™ Technologies |
| N2 Supplement-A | 7152 | STEMCELL™ Technologies |

**Table S11:**
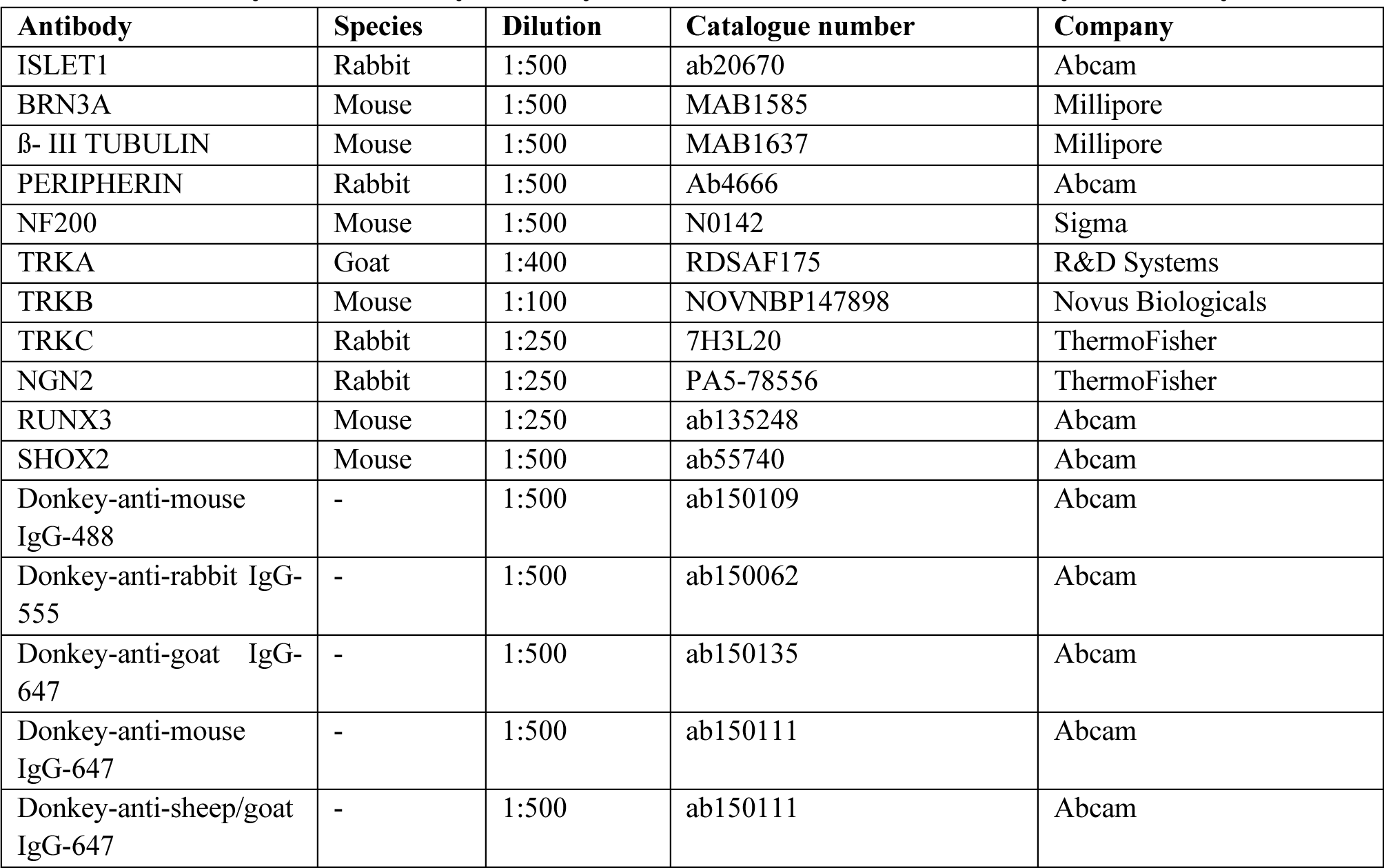
Primary and secondary antibody details and dilutions for immunocytochemistry.

**Table S12:** Primary and secondary antibody details for western blotting.

| Antibody name | Host | Species reactivity | Dilution | Catalogue number | Company |
| --- | --- | --- | --- | --- | --- |
| NGN2 | Rabbit | Human, Mouse, Rat | 1:3000 | PA5-78556 | ThermoFisher |
| RUNX3 | Mouse | Human, mouse | 1:3000 | ab135248 | Abcam |
| SHOX2 | Mouse | Human, rat | 1:3000 | ab55740 | Abcam |
| GAPDH | Mouse | Human | 1:20,000 | G8795 | Sigma |
| GAPDH | Rabbit | Human | 1:20,000 | G9545 | Sigma |
| Goat Anti-Mouse IgG H&L (HRP) | - | - | 1:10,000 | ab97023 | Abcam |
| Goat Anti-Rabbit IgG Antibody,<br>(H+L) HRP conjugate | - | - | 1:10,000 | ap307p | Merk Millipore |

## Supplementary Figures

**Supplementary Figure 1:**
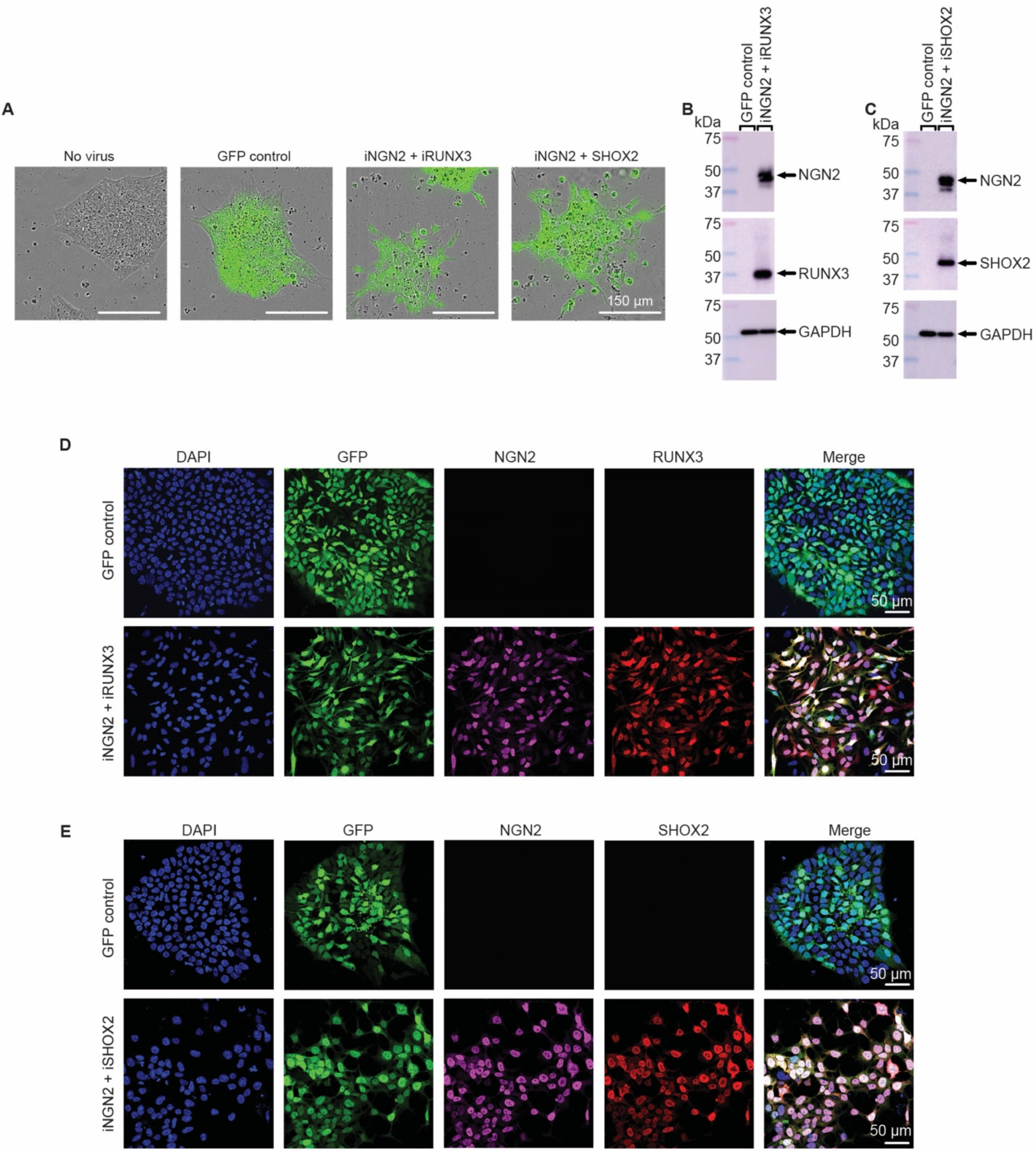
Validation of GFP, NGN2 and either RUNX3 or SHOX2 expression in hPSCs. hPSCs were transduced with lentiviral viral particles containing the reverse tetracycline transactivator (rTTA) and either pLV-TetO-eGFP-PuroR (GFP control), pLV-TetO-hNGN2-hRUNX3-GFP-PuroR (iNGN2 + iRUNX3) or pLV-TetO-hNGN2-hSHOX2-GFP-PuroR (iNGN2 + iSHOX2). Viral particles were removed and doxycycline was administered 24 h following transduction. hPSCs were harvested for Western blotting or immunocytochemistry after 96 h of doxycycline administration. (A) Representative images of transduced cultures that confirm the expression of GFP after 96 h of doxycycline incubation. Western blots of hPSC protein lysates probed for (B) NGN2 and RUNX3 or (C) NGN2 and SHOX2 protein expression. (D) Representative immunocytochemistry images showing the cellular co-localisation of GFP (*green*) with NGN2 (*magenta*) and RUNX3 (*red*) or (E) with NGN2 (*magenta*) and SHOX2 (*red*). Nuclei are shown in blue. Scale bars = 50 μm.

**Supplementary Figure 2:**
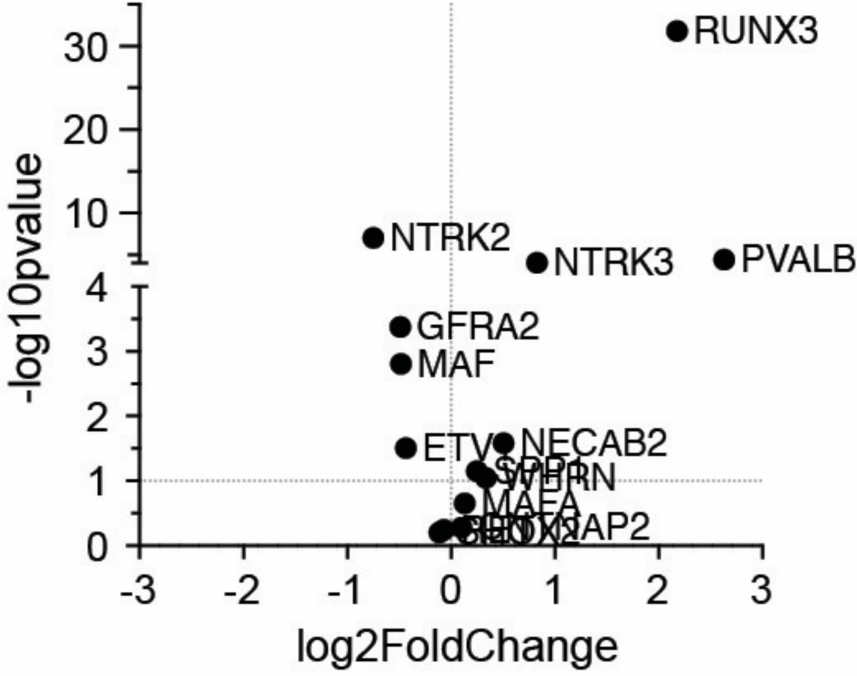
Enrichment of PN and LTMR markers in the iPN and iLTMR cultures, respectively. Differential gene expression plot comparing the expression of transcripts associated with PNs and LTMRs in PNs (*right*) versus LTMRs (*left*) as determined by bulk RNA sequencing. n = 3 biological replicates.

**Supplementary Figure 3:**
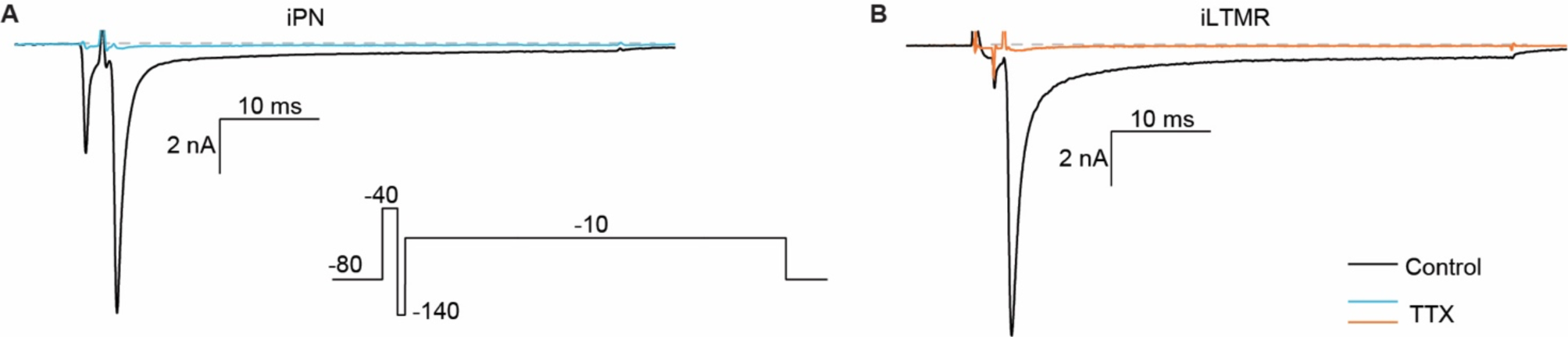
TTX inhibition of iPN and iLTMR I_Nav_. Representative Nav currents in (A) iPN and (B) iLTMRs in the absence (control) and presence of TTX (300 nM). Inset: stimulus: 50 ms, −10 mV, Vh −80 mV, 0.1Hz). Scale bars: 2 nA, 10 ms. Numeric data are included in Table S3.

**Supplementary Figure 4:**
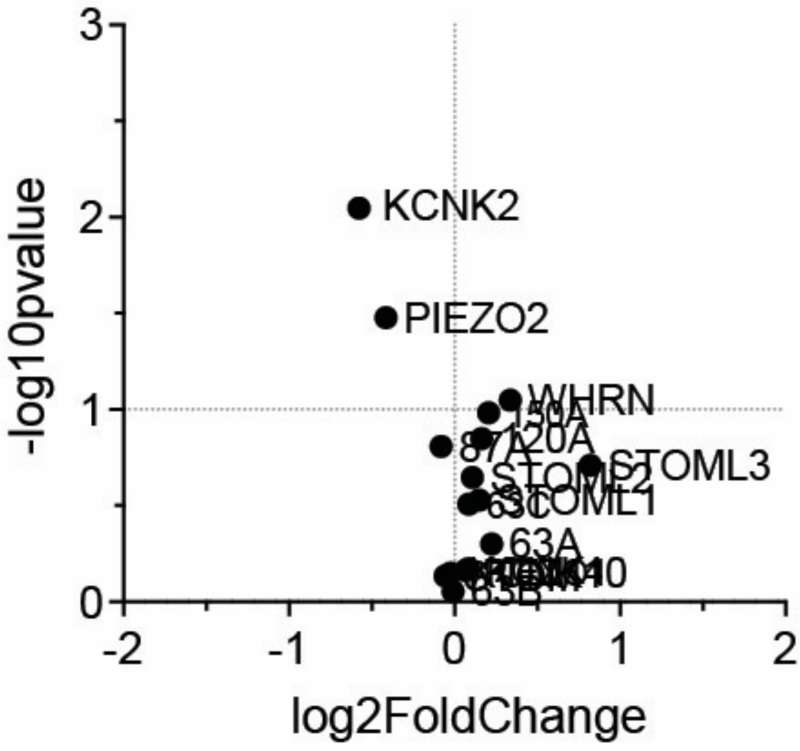
Differential gene expression plot comparing mechanosensitivity-associated transcripts between iPNs and iLTMRs. The major gene transcripts associated with the detection of mechanical sensations by comparing the iPN expression (*right*) against the iLTMR expression (*left*), from bulk RNA sequencing. n = 3 biological replicates.

**Supplementary Figure 5:**
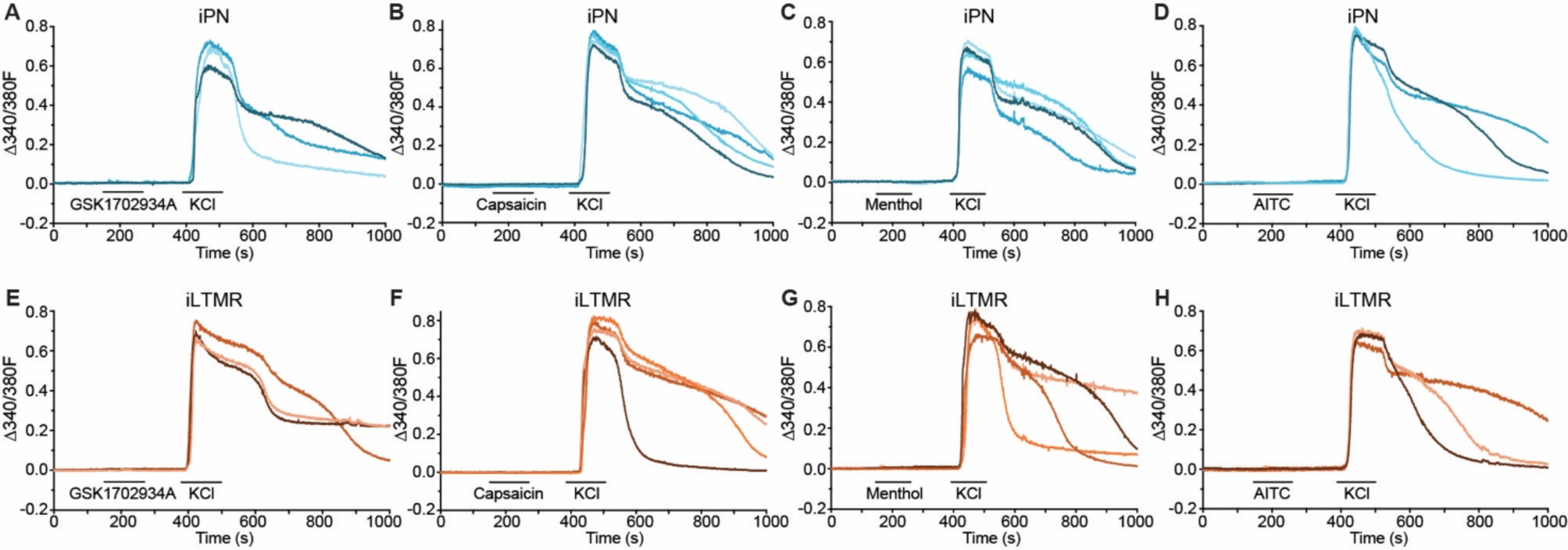
Absence of Nociceptive Responses in iPNs and iLTMRs. Representative live cell Fura-2 calcium imaging traces of iPNs (*top*) in the presence of (A) 1 µM GSK1702934A (TRPC3/6 activator), (B) 1 µM capsaicin (TRPV1 activator), (C) 250 µM menthol (TRPM8 activator) and (D) 100 µM AITC (TRPA1 activator). Representative live cell Fura-2 calcium imaging traces of iLTMRs (*bottom*) in the presence of (E) 1 µM GSK1702934A, (F) 1 µM capsaicin, (G) 250 µM menthol and (H) 100 µM AITC. Traces represent the change in the 340/380 fluorescence ratio (1′340/380F) (imaged every 0.7 s) from baseline, of 3 – 4 individual neurons perfused (1 mL/min) with the selected agonist followed by 60 mM KCl with CBS washes before and after treatments. n > 100 neurons per agonist across 3 biological replicates. Numeric data are included in Table S4.

**Supplementary Figure 6:**
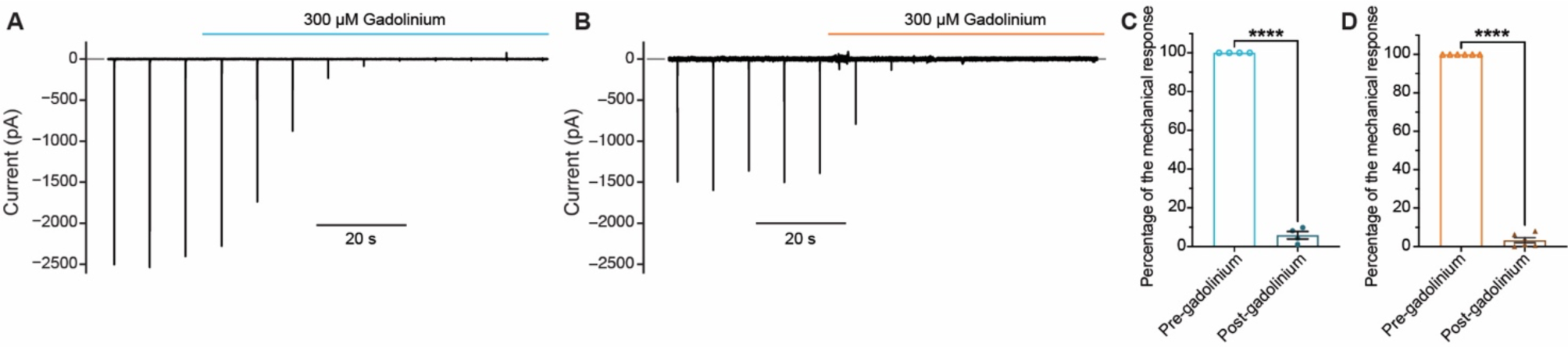
Gadolinium inhibits MA currents in iPNs and iLTMRs. Representative whole-cell voltage-clamp recording of an (A) iPN and (B) iLTMR neuron mechanically stimulated by repetitive 0.5 μm (100 ms duration) membrane probe indentations, followed by direct bath addition of 300 μM gadolinium in extracellular buffer. Normalised percentages of the mechanical response, relative to the average mechanical response before and after gadolinium application, in (C) iPN and (D) iLTMR. Unpaired t-test, ****p < 0.0001. n = 3 – 4 biological replicates, n = 4 – 6 neurons in total. Data shown represents the mean ± SEM.

**Supplementary Figure 7:**
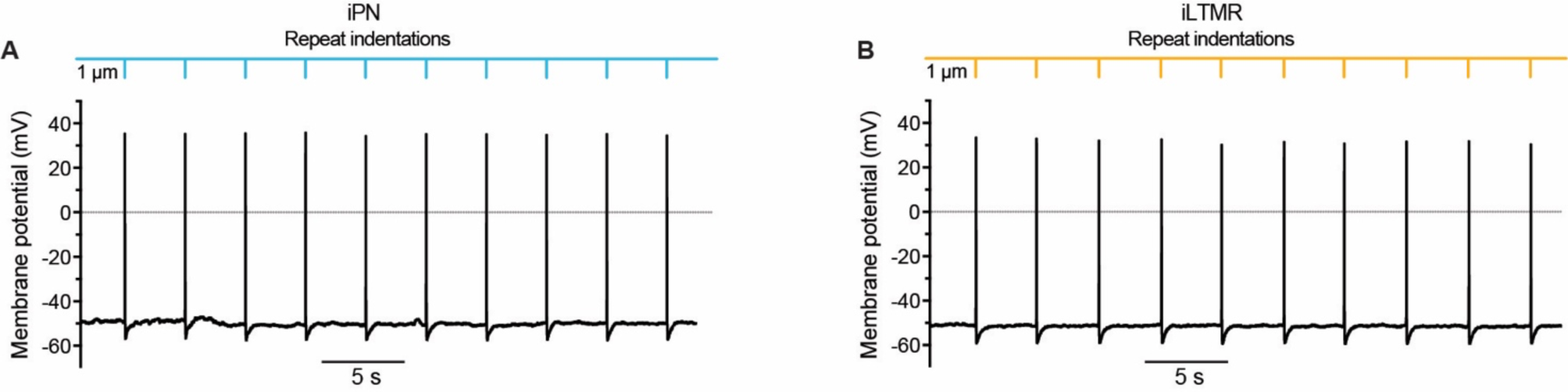
iPNs and iLTMRs fire stereotypical action potentials in response to mechanical stimulation. Representative whole-cell current-clamp recording of an (A) iPN and (B) iLTMR neuron mechanically stimulated by repeated 1 µm membrane probe indentations (100 ms duration).

**Supplementary Figure 8:**
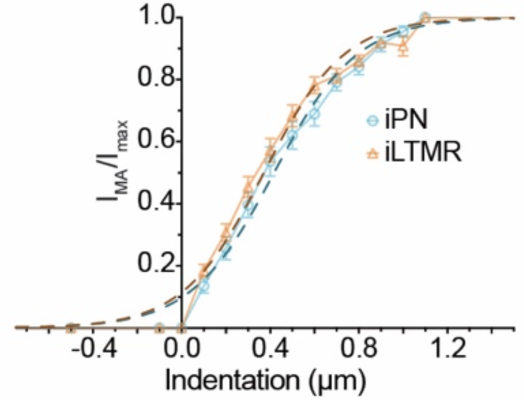
I_MA_ activation vs indentation plot. The I_MA_ (from Fig. 4) normalised to the I_MAX_ and plotted against increasing membrane probe indentation. n = 25 – 33 neurons across n = 5 – 8 biological replicates.

**Supplementary Figure 9:**
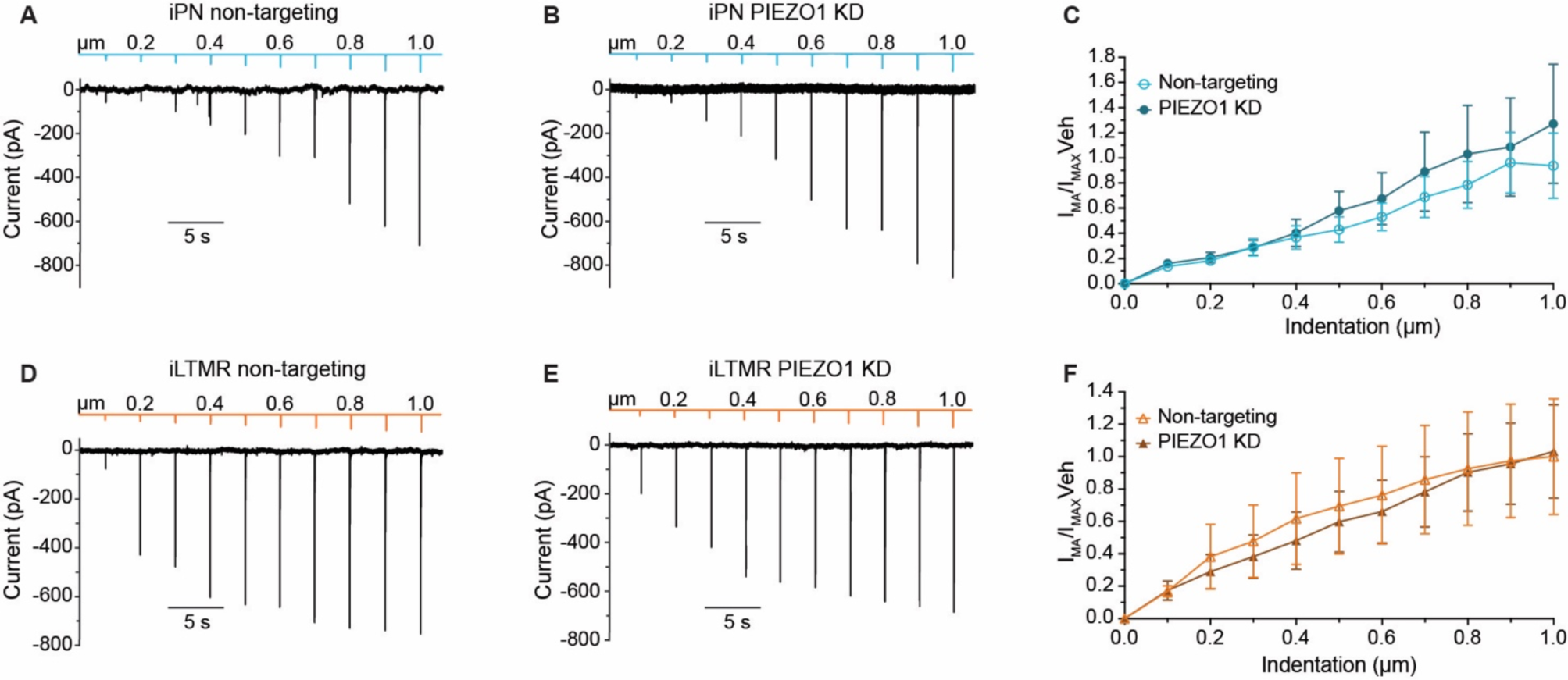
PIEZO1 does not contribute to the MA current in the iPNs and iLTMRs. iPNs and iLTMRs under whole-cell voltage-clamp conditions with increasing 0.1 μm increments of membrane probe indentation (from 0 – 1 μm, Δ 0.1 μm). Representative traces of an iPN transfected with (A) non-targeting siRNA or (B) PIEZO1 siRNA, for 96 h. (C) The current density of iPNs in response to indentation normalised to the average non-targeting siRNA response at 1 μm indentation. Representative trace of an iLTMR transfected with (D) non-targeting siRNA or (E) PIEZO1 siRNA, for 96 h. (F) The current density of iLTMRs in response to indentation normalised to the average non-targeting siRNA response at 1 μm indentation. Data represent mean ±SEM, n = 15 – 16 neurons across 3 independent siRNA transfections.

## Notes

### Competing Interest Statement

The authors have declared no competing interest.

## References

1. Handler, A., and Ginty, D.D. (2021). The mechanosensory neurons of touch and their mechanisms of activation. Nat Rev Neurosci 22, 521–537. 10.1038/s41583-021-00489-x.

2. Hao, J., and Delmas, P. (2010). Multiple desensitization mechanisms of mechanotransducer channels shape firing of mechanosensory neurons. J Neurosci 30, 13384–13395. 10.1523/JNEUROSCI.2926-10.2010.

3. Parpaite, T., Brosse, L., Séjourné, N., Laur, A., Mechioukhi, Y., Delmas, P., and Coste, B. (2021). Patch-seq of mouse DRG neurons reveals candidate genes for specific mechanosensory functions. Cell Rep 37, 109914. 10.1016/j.celrep.2021.109914.

4. Zheng, Y., Liu, P., Bai, L., Trimmer, J.S., Bean, B.P., and Ginty, D.D. (2019). Deep Sequencing of Somatosensory Neurons Reveals Molecular Determinants of Intrinsic Physiological Properties. Neuron 103, 598–616.e7. 10.1016/j.neuron.2019.05.039.

5. Li, C.-L., Li, K.-C., Wu, D., Chen, Y., Luo, H., Zhao, J.-R., Wang, S.-S., Sun, M.-M., Lu, Y.-J., Zhong, Y.-Q., et al. (2016). Somatosensory neuron types identified by high-coverage single-cell RNA-sequencing and functional heterogeneity. Cell Res 26, 967. 10.1038/cr.2016.90.

6. Usoskin, D., Furlan, A., Islam, S., Abdo, H., Lönnerberg, P., Lou, D., Hjerling-Leffler, J., Haeggström, J., Kharchenko, O., Kharchenko, P.V., et al. (2015). Unbiased classification of sensory neuron types by large-scale single-cell RNA sequencing. Nat Neurosci 18, 145–153. 10.1038/nn.3881.

7. Wu, H., Petitpré, C., Fontanet, P., Sharma, A., Bellardita, C., Quadros, R.M., Jannig, P.R., Wang, Y., Heimel, J.A., Cheung, K.K.Y., et al. (2021). Distinct subtypes of proprioceptive dorsal root ganglion neurons regulate adaptive proprioception in mice. Nat Commun 12, 1026. 10.1038/s41467-021-21173-9.

8. Chang, W., Berta, T., Kim, Y.H., Lee, S., Lee, S.-Y., and Ji, R.-R. (2018). Expression and Role of Voltage-Gated Sodium Channels in Human Dorsal Root Ganglion Neurons with Special Focus on Nav1.7, Species Differences, and Regulation by Paclitaxel. Neurosci Bull 34, 4–12. 10.1007/s12264-017-0132-3.

9. Schwaid, A.G., Krasowka-Zoladek, A., Chi, A., and Cornella-Taracido, I. (2018). Comparison of the Rat and Human Dorsal Root Ganglion Proteome. Sci Rep 8, 13469. 10.1038/s41598-018-31189-9.

10. Rostock, C., Schrenk-Siemens, K., Pohle, J., and Siemens, J. (2018). Human vs. Mouse Nociceptors -Similarities and Differences. Neuroscience 387, 13–27. 10.1016/j.neuroscience.2017.11.047.

11. Jung, M., Dourado, M., Maksymetz, J., Jacobson, A., Laufer, B.I., Baca, M., Foreman, O., Hackos, D.H., Riol-Blanco, L., and Kaminker, J.S. (2023). Cross-species transcriptomic atlas of dorsal root ganglia reveals species-specific programs for sensory function. Nat Commun 14, 366. 10.1038/s41467-023-36014-0.

12. Hulme, A.J., Maksour, S., St-Clair Glover, M., Miellet, S., and Dottori, M. (2022). Making neurons, made easy: The use of Neurogenin-2 in neuronal differentiation. Stem Cell Reports 17, 14–34. 10.1016/j.stemcr.2021.11.015.

13. Masserdotti, G., Gascón, S., and Götz, M. (2016). Direct neuronal reprogramming: learning from and for development. Development 143, 2494–2510. 10.1242/dev.092163.

14. Nickolls, A.R., Lee, M.M., Espinoza, D.F., Szczot, M., Lam, R.M., Wang, Q., Beers, J., Zou, J., Nguyen, M.Q., Solinski, H.J., et al. (2020). Transcriptional Programming of Human Mechanosensory Neuron Subtypes from Pluripotent Stem Cells. Cell Rep 30, 932–946.e7. 10.1016/j.celrep.2019.12.062.

15. Schrenk-Siemens, K., Wende, H., Prato, V., Song, K., Rostock, C., Loewer, A., Utikal, J., Lewin, G.R., Lechner, S.G., and Siemens, J. (2015). PIEZO2 is required for mechanotransduction in human stem cell-derived touch receptors. Nat Neurosci 18, 10–16. 10.1038/nn.3894.

16. Schrenk-Siemens, K., Pohle, J., Rostock, C., Abd El Hay, M., Lam, R.M., Szczot, M., Lu, S., Chesler, A.T., and Siemens, J. (2022). Human Stem Cell-Derived TRPV1-Positive Sensory Neurons: A New Tool to Study Mechanisms of Sensitization. Cells 11, 2905. 10.3390/cells11182905.

17. Blanchard, J.W., Eade, K.T., Szűcs, A., Lo Sardo, V., Tsunemoto, R.K., Williams, D., Sanna, P.P., and Baldwin, K.K. (2015). Selective conversion of fibroblasts into peripheral sensory neurons. Nat Neurosci 18, 25–35. 10.1038/nn.3887.

18. Hulme, A.J., McArthur, J.R., Maksour, S., Miellet, S., Ooi, L., Adams, D.J., Finol-Urdaneta, R.K., and Dottori, M. (2020). Molecular and Functional Characterization of Neurogenin-2 Induced Human Sensory Neurons. Front Cell Neurosci 14, 600895. 10.3389/fncel.2020.600895.

19. Plumbly, W., Patikas, N., Field, S.F., Foskolou, S., and Metzakopian, E. (2022). Derivation of nociceptive sensory neurons from hiPSCs with early patterning and temporally controlled NEUROG2 overexpression. Cell Rep Methods 2, 100341. 10.1016/j.crmeth.2022.100341.

20. Abdo, H., Li, L., Lallemend, F., Bachy, I., Xu, X.-J., Rice, F.L., and Ernfors, P. (2011). Dependence on the transcription factor Shox2 for specification of sensory neurons conveying discriminative touch. Eur J Neurosci 34, 1529–1541. 10.1111/j.1460-9568.2011.07883.x.

21. Inoue, K., Ito, K., Osato, M., Lee, B., Bae, S.-C., and Ito, Y. (2007). The transcription factor Runx3 represses the neurotrophin receptor TrkB during lineage commitment of dorsal root ganglion neurons. J Biol Chem 282, 24175–24184. 10.1074/jbc.M703746200.

22. Kramer, I., Sigrist, M., de Nooij, J.C., Taniuchi, I., Jessell, T.M., and Arber, S. (2006). A role for Runx transcription factor signaling in dorsal root ganglion sensory neuron diversification. Neuron 49, 379–393. 10.1016/j.neuron.2006.01.008.

23. Nakamura, S., Senzaki, K., Yoshikawa, M., Nishimura, M., Inoue, K.-I., Ito, Y., Ozaki, S., and Shiga, T. (2008). Dynamic regulation of the expression of neurotrophin receptors by Runx3. Development 135, 1703–1711. 10.1242/dev.015248.

24. Ogihara, Y., Masuda, T., Ozaki, S., Yoshikawa, M., and Shiga, T. (2016). Runx3-regulated expression of two Ntrk3 transcript variants in dorsal root ganglion neurons. Dev Neurobiol 76, 313–322. 10.1002/dneu.22316.

25. Scott, A., Hasegawa, H., Sakurai, K., Yaron, A., Cobb, J., and Wang, F. (2011). Transcription factor short stature homeobox 2 is required for proper development of tropomyosin-related kinase B-expressing mechanosensory neurons. J Neurosci 31, 6741–6749. 10.1523/JNEUROSCI.5883-10.2011.

26. Alshawaf, A.J., Viventi, S., Qiu, W., D’Abaco, G., Nayagam, B., Erlichster, M., Chana, G., Everall, I., Ivanusic, J., Skafidas, E., et al. (2018). Phenotypic and Functional Characterization of Peripheral Sensory Neurons derived from Human Embryonic Stem Cells. Sci Rep 8, 603. 10.1038/s41598-017-19093-0.

27. Espino, C.M., Lewis, C.M., Ortiz, S., Dalal, M.S., Garlapalli, S., Wells, K.M., O’Neil, D.A., Wilkinson, K.A., and Griffith, T.N. (2022). Na_V_1.1 is essential for proprioceptive signaling and motor behaviors. Elife 11, e79917. 10.7554/eLife.79917.

28. Ranade, S.S., Woo, S.-H., Dubin, A.E., Moshourab, R.A., Wetzel, C., Petrus, M., Mathur, J., Bégay, V., Coste, B., Mainquist, J., et al. (2014). Piezo2 is the major transducer of mechanical forces for touch sensation in mice. Nature 516, 121–125. 10.1038/nature13980.

29. Woo, S.-H., Lukacs, V., de Nooij, J.C., Zaytseva, D., Criddle, C.R., Francisco, A., Jessell, T.M., Wilkinson, K.A., and Patapoutian, A. (2015). Piezo2 is the principal mechanotransduction channel for proprioception. Nat Neurosci 18, 1756–1762. 10.1038/nn.4162.

30. Lengyel, M., Enyedi, P., and Czirják, G. (2021). Negative Influence by the Force: Mechanically Induced Hyperpolarization via K2P Background Potassium Channels. International Journal of Molecular Sciences 22, 9062. 10.3390/ijms22169062.

31. Moehring, F., Halder, P., Seal, R.P., and Stucky, C.L. (2018). Uncovering the Cells and Circuits of Touch in Normal and Pathological Settings. Neuron 100, 349–360. 10.1016/j.neuron.2018.10.019.

32. Patel, A.J., Honoré, E., Maingret, F., Lesage, F., Fink, M., Duprat, F., and Lazdunski, M. (1998). A mammalian two pore domain mechano-gated S-like K+ channel. EMBO J 17, 4283–4290. 10.1093/emboj/17.15.4283.

33. Wetzel, C., Pifferi, S., Picci, C., Gök, C., Hoffmann, D., Bali, K.K., Lampe, A., Lapatsina, L., Fleischer, R., Smith, E.S.J., et al. (2017). Small-molecule inhibition of STOML3 oligomerization reverses pathological mechanical hypersensitivity. Nat Neurosci 20, 209–218. 10.1038/nn.4454.

34. Poole, K., Herget, R., Lapatsina, L., Ngo, H.-D., and Lewin, G.R. (2014). Tuning Piezo ion channels to detect molecular-scale movements relevant for fine touch. Nat Commun 5, 3520. 10.1038/ncomms4520.

35. de Nooij, J.C., Simon, C.M., Simon, A., Doobar, S., Steel, K.P., Banks, R.W., Mentis, G.Z., Bewick, G.S., and Jessell, T.M. (2015). The PDZ-domain protein Whirlin facilitates mechanosensory signaling in mammalian proprioceptors. J Neurosci 35, 3073–3084. 10.1523/JNEUROSCI.3699-14.2015.

36. Friedrich, O., Merten, A.-L., Schneidereit, D., Guo, Y., Schürmann, S., and Martinac, B. (2019). Stretch in Focus: 2D Inplane Cell Stretch Systems for Studies of Cardiac Mechano-Signaling. Front Bioeng Biotechnol 7, 55. 10.3389/fbioe.2019.00055.

37. Schürmann, S., Wagner, S., Herlitze, S., Fischer, C., Gumbrecht, S., Wirth-Hücking, A., Prölß, G., Lautscham, L.A., Fabry, B., Goldmann, W.H., et al. (2016). The IsoStretcher: An isotropic cell stretch device to study mechanical biosensor pathways in living cells. Biosens Bioelectron 81, 363–372. 10.1016/j.bios.2016.03.015.

38. Guo, Y., Merten, A.-L., Schöler, U., Yu, Z.-Y., Cvetkovska, J., Fatkin, D., Feneley, M.P., Martinac, B., and Friedrich, O. (2021). In vitro cell stretching technology (IsoStretcher) as an approach to unravel Piezo1-mediated cardiac mechanotransduction. Prog Biophys Mol Biol 159, 22–33. 10.1016/j.pbiomolbio.2020.07.003.

39. Bhattacharya, M.R.C., Bautista, D.M., Wu, K., Haeberle, H., Lumpkin, E.A., and Julius, D. (2008). Radial stretch reveals distinct populations of mechanosensitive mammalian somatosensory neurons. Proc Natl Acad Sci U S A 105, 20015–20020. 10.1073/pnas.0810801105.

40. Ermakov, Y.A., Kamaraju, K., Sengupta, K., and Sukharev, S. (2010). Gadolinium ions block mechanosensitive channels by altering the packing and lateral pressure of anionic lipids. Biophys J 98, 1018–1027. 10.1016/j.bpj.2009.11.044.

41. Rugiero, F., Drew, L.J., and Wood, J.N. (2010). Kinetic properties of mechanically activated currents in spinal sensory neurons. J Physiol 588, 301–314. 10.1113/jphysiol.2009.182360.

42. Hill, R.Z., Loud, M.C., Dubin, A.E., Peet, B., and Patapoutian, A. (2022). PIEZO1 transduces mechanical itch in mice. Nature 607, 104–110. 10.1038/s41586-022-04860-5.

43. Szczot, M., Pogorzala, L.A., Solinski, H.J., Young, L., Yee, P., Le Pichon, C.E., Chesler, A.T., and Hoon, M.A. (2017). Cell-Type-Specific Splicing of Piezo2 Regulates Mechanotransduction. Cell Rep 21, 2760–2771. 10.1016/j.celrep.2017.11.035.

44. Vega-Bermudez, F., and Johnson, K.O. (1999). SA1 and RA receptive fields, response variability, and population responses mapped with a probe array. J Neurophysiol 81, 2701–2710. 10.1152/jn.1999.81.6.2701.

45. Hong, G.-S., Lee, B., Wee, J., Chun, H., Kim, H., Jung, J., Cha, J.Y., Riew, T.-R., Kim, G.H., Kim, I.-B., et al. (2016). Tentonin 3/TMEM150c Confers Distinct Mechanosensitive Currents in Dorsal-Root Ganglion Neurons with Proprioceptive Function. Neuron 91, 107–118. 10.1016/j.neuron.2016.05.029.

46. Hunt, C.C., and Ottoson, D. (1975). Impulse activity and receptor potential of primary and secondary endings of isolated mammalian muscle spindles. J Physiol 252, 259–281. 10.1113/jphysiol.1975.sp011143.

47. Abraira, V.E., and Ginty, D.D. (2013). The sensory neurons of touch. Neuron 79, 618–639. 10.1016/j.neuron.2013.07.051.

48. Johnson, K.O. (2001). The roles and functions of cutaneous mechanoreceptors. Curr Opin Neurobiol 11, 455–461. 10.1016/s0959-4388(00)00234-8.

49. Johnson, K.O., Yoshioka, T., and Vega-Bermudez, F. (2000). Tactile functions of mechanoreceptive afferents innervating the hand. J Clin Neurophysiol 17, 539–558. 10.1097/00004691-200011000-00002.

50. François, A., Schüetter, N., Laffray, S., Sanguesa, J., Pizzoccaro, A., Dubel, S., Mantilleri, A., Nargeot, J., Noël, J., Wood, J.N., et al. (2015). The Low-Threshold Calcium Channel Cav3.2 Determines Low-Threshold Mechanoreceptor Function. Cell Rep 10, 370–382. 10.1016/j.celrep.2014.12.042.

51. Lipovsek, M., Bardy, C., Cadwell, C.R., Hadley, K., Kobak, D., and Tripathy, S.J. (2021). Patch-seq: Past, Present, and Future. J Neurosci 41, 937–946. 10.1523/JNEUROSCI.1653-20.2020.

52. Handler, A., Zhang, Q., Pang, S., Nguyen, T.M., Iskols, M., Nolan-Tamariz, M., Cattel, S., Plumb, R., Sanchez, B., Ashjian, K., et al. (2023). Three-dimensional reconstructions of mechanosensory end organs suggest a unifying mechanism underlying dynamic, light touch. Neuron 111, 3211–3229.e9. 10.1016/j.neuron.2023.08.023.

53. Mamiya, A., Sustar, A., Siwanowicz, I., Qi, Y., Lu, T.-C., Gurung, P., Chen, C., Phelps, J.S., Kuan, A.T., Pacureanu, A., et al. (2023). Biomechanical origins of proprioceptor feature selectivity and topographic maps in the Drosophila leg. Neuron 111, 3230–3243.e14. 10.1016/j.neuron.2023.07.009.

54. Nikolaev, Y.A., Feketa, V.V., Anderson, E.O., Schneider, E.R., Gracheva, E.O., and Bagriantsev, S.N. (2020). Lamellar cells in Pacinian and Meissner corpuscles are touch sensors. Sci Adv 6, eabe6393. 10.1126/sciadv.abe6393.

55. Ziolkowski, L.H., Gracheva, E.O., and Bagriantsev, S.N. (2023). Mechanotransduction events at the physiological site of touch detection. eLife 12, e84179. 10.7554/eLife.84179.

56. Chakrabarti, S., Klich, J.D., Khallaf, M.A., Hulme, A.J., Sánchez-Carranza, O., Baran, Z.M., Rossi, A., Huang, A.T.-L., Pohl, T., Fleischer, R., et al. (2024). Touch sensation requires the mechanically gated ion channel ELKIN1. Science 383, 992–998. 10.1126/science.adl0495.

57. Romero, L.O., Massey, A.E., Mata-Daboin, A.D., Sierra-Valdez, F.J., Chauhan, S.C., Cordero-Morales, J.F., and Vásquez, V. (2019). Dietary fatty acids fine-tune Piezo1 mechanical response. Nat Commun 10, 1200. 10.1038/s41467-019-09055-7.

58. Vásquez, V., Krieg, M., Lockhead, D., and Goodman, M.B. (2014). Phospholipids that contain polyunsaturated fatty acids enhance neuronal cell mechanics and touch sensation. Cell Rep 6, 70–80. 10.1016/j.celrep.2013.12.012.

59. Koeppen, A.H. (2011). Friedreich’s ataxia: pathology, pathogenesis, and molecular genetics. J Neurol Sci 303, 1–12. 10.1016/j.jns.2011.01.010.

60. Pandolfo, M. (2009). Friedreich ataxia: the clinical picture. J Neurol 256 *Suppl 1*, 3–8. 10.1007/s00415-009-1002-3.

61. Dhandapani, R., Arokiaraj, C.M., Taberner, F.J., Pacifico, P., Raja, S., Nocchi, L., Portulano, C., Franciosa, F., Maffei, M., Hussain, A.F., et al. (2018). Control of mechanical pain hypersensitivity in mice through ligand-targeted photoablation of TrkB-positive sensory neurons. Nat Commun 9, 1640. 10.1038/s41467-018-04049-3.

62. Dothel, G., Barbaro, M.R., Boudin, H., Vasina, V., Cremon, C., Gargano, L., Bellacosa, L., De Giorgio, R., Le Berre-Scoul, C., Aubert, P., et al. (2015). Nerve fiber outgrowth is increased in the intestinal mucosa of patients with irritable bowel syndrome. Gastroenterology 148, 1002–1011.e4. 10.1053/j.gastro.2015.01.042.

63. Orefice, L.L. (2020). Peripheral Somatosensory Neuron Dysfunction: Emerging Roles in Autism Spectrum Disorders. Neuroscience 445, 120–129. 10.1016/j.neuroscience.2020.01.039.

64. Schaffler, M.D., Middleton, L.J., and Abdus-Saboor, I. (2019). Mechanisms of Tactile Sensory Phenotypes in Autism: Current Understanding and Future Directions for Research. Curr Psychiatry Rep 21, 134. 10.1007/s11920-019-1122-0.

65. Wang, R., and Lewin, G.R. (2011). The Cav3.2 T-type calcium channel regulates temporal coding in mouse mechanoreceptors. J Physiol 589, 2229–2243. 10.1113/jphysiol.2010.203463.

66. Zhang, Y., Parmigiani, G., and Johnson, W.E. (2020). ComBat-seq: batch effect adjustment for RNA-seq count data. NAR Genom Bioinform 2, lqaa078. 10.1093/nargab/lqaa078.

67. Love, M.I., Huber, W., and Anders, S. (2014). Moderated estimation of fold change and dispersion for RNA-seq data with DESeq2. Genome Biol 15, 550. 10.1186/s13059-014-0550-8.

